# Increased Reproducibility of Brain Organoids through Controlled Fluid Dynamics

**DOI:** 10.1101/2025.03.05.641438

**Authors:** Giuseppe Aiello, Mohamed Nemir, Barbora Vidimova, Cindy Ramel, Joanna Viguie, Arianna Ravera, Krzysztof Wrzesinski, Claudia Bagni

## Abstract

Brain organoids are a promising model for studying human neurodevelopment and disease. Despite the potential, their 3D structure often exhibits high variability during differentiation across batches and cell lines, presenting a significant challenge for biomedical applications. During their development, organoids are exposed to fluid flow shear stress (fFSS) generated by the flow of culture media over the developing tissue. This stress is thought to disrupt cellular integrity and morphogenesis, leading to variation in organoids architecture, ultimately affecting reproducibility. Understanding the interplay between tissue morphology, cell identity and organoid developmental stage is therefore essential for advancing the use of brain organoids. Here, we demonstrate that reducing fFSS, by employing a vertically rotating chamber during neuronal induction, a critical phase for organoid morphogenesis, along with an extended human induced pluripotent stem cell (hiPSC) aggregation phase to minimize fusions, significantly improves the reproducibility of brain organoids. Remarkably, reducing fFSS minimized morphological structure variation and preserved transcriptional signature fidelity across differentiation batches and cell lines. This approach could enhance the reliability of brain organoid models, with important implications for neurodevelopmental research and preclinical studies.

## INTRODUCTION

Brain organoids have become invaluable tools for advancing our understanding of human neurodevelopment, modelling neurological disorders and exploring personalized medicine. These models, compared to classic differentiation models in monolayer cultures, have shown higher complexity and similarity to the developing human brain, offering an insight into processes that were previously inaccessible (Antonica *et al*, 2022; Camp *et al*, 2015; Eichmuller & Knoblich, 2022; Kelley & Pasca, 2022; Kim *et al*, 2020; Lancaster *et al*, 2013; Luo *et al*, 2016; Pellegrini *et al*, 2020; Qian *et al*, 2019).

A key strength of brain organoids lies in their ability to mimic the intricate relationship between tissue structure and gene expression, where precise morphological organization underpins the tissue’s functional complexity (Camp *et al*., 2015; Fleck *et al*, 2023; Luo *et al*., 2016; Quadrato *et al*, 2017; Warmflash *et al*, 2014). However, brain organoid technology faces a critical barrier to widespread adoption related to reproducibility (Di Lullo & Kriegstein, 2017; Qian *et al*., 2019). Variability in tissue morphology, cellular composition and differentiation outcomes across batches and cell lines limits their reliability. This inconsistency poses significant challenges for comparative studies and high-throughput applications, reducing their utility in both basic and translational research (Bhaduri *et al*, 2020; Lancaster *et al*, 2017; Messana *et al*, 2008; Velasco *et al*, 2019).

Tissue morphology is increasingly recognized as a critical determinant of organoid development, and recent studies have highlighted its importance in guiding the temporal and spatial aspects of this process (Gjorevski *et al*, 2022; Lancaster *et al*., 2017). Morphological inconsistencies not only affect the structural fidelity of organoids but also alter the temporal program of differentiation, impacting regional identity, cellular organization and functional properties (Chiaradia *et al*, 2023; Jain *et al*, 2023; Martins-Costa *et al*, 2023).

The use of extra-cellular matrix (ECM) substrates, such as solubilized basement membrane Matrigel, is a widely adopted culturing strategy. Nonetheless, it is known to influence organoid morphology and introduce variability in differentiation outcomes, therefore having an impact on reproducibility across experiments (Jain *et al*., 2023; Long *et al*, 2018; Martins-Costa *et al*., 2023).

These insights underscore the need for methods that standardize the morphological aspects of brain organoid development, as irregularities in tissue structure can propagate through later stages of development, contributing to variable phenotypical outcomes. While much attention has been given to intrinsic factors, such as genetic variability between cell lines, the influence of extrinsic biomechanical factors is often underestimated.

In the developing embryo, mechanical forces play a fundamental role in shaping tissues and guiding differentiation: contractility and tension create a mechano-transduction feedback loop that regulates morphogenesis and patterning, influencing gene expression through pathways such as SRC-dependent β-catenin signalling (Caldarelli *et al*, 2024). This highlights the intricate crosstalk between mechanical forces and molecular pathways as key regulators of development.

Brain organoids, despite lacking maternal cues and extraembryonic tissues, are highly sensitive to external mechanical perturbations, particularly during the earliest stages of differentiation. The initial aggregation and the subsequent neuronal induction phases are critical windows where tissue identity and patterning are established. During these stages, cells undergo self-organization, lineage specification and neuroepithelium formation, processes that depend on finely balanced mechanical cues (Andrews & Nowakowski, 2019; Renner *et al*, 2017).

In the development of brain organoids, extrinsic force variation can disrupt tissue integrity and morphogenesis, leading to organoids that differ in shape and functionality (Goto-Silva *et al*, 2019; Ishihara *et al*, 2023; Saglam-Metiner *et al*, 2023). One such factor is fluid flow shear stress (fFSS), which arises from the flow of culture media in dynamic systems commonly used for organoid culture (Dahl-Jensen & Grapin-Botton, 2017; Goto-Silva *et al*., 2019; Hinton *et al*, 2022). These systems, including plates, bioreactors and spinning bioreactors, are designed to improve nutrient and oxygen distribution, promoting cell survival and growth. However, the trade-off is the generation of fFSS, a mechanical force that can disrupt cellular integrity and tissue organization (Croughan & Wang, 1991; Goto-Silva *et al*., 2019; Suong *et al*, 2021). Prolonged exposure to fFSS during critical stages of organoid development has been shown to interfere with proper morphogenesis, leading to inconsistencies in tissue architecture that ultimately affect reproducibility (Dardik *et al*, 2005; Gareau *et al*, 2014; Ismadi *et al*, 2014; Saglam-Metiner *et al*., 2023).

Additionally, cellular metabolism is increasingly recognized as a key driver of neurodevelopment, influencing energy states, signalling pathways and overall tissue health (Badal *et al*, 2019; Iwata *et al*, 2023; Iwata *et al*, 2020; Khacho & Slack, 2018). Metabolic heterogeneity in brain organoids is tightly linked to differentiation outcomes, with variations in oxidative phosphorylation (OXPHOS) and glycolysis influencing progenitor maintenance and neuronal specification (Øhlenschlæger *et al*, 2023). Nonetheless, it remains unclear whether changes in metabolic states across conditions in organoids are linked to variability in cellular composition or genuinely driven by biological differences.

Here we demonstrate that reducing fFSS during neuronal induction, a critical phase of cortical patterning, significantly improves the morphological reproducibility of brain organoids. By employing a rotating chamber (RC) apparatus, designed to minimize fFSS, and using an ECM-free approach for greater control over differentiation conditions, we achieved consistency of morphological features and transcriptional fidelity across organoid batches and cell lines. Furthermore, our transcriptional analysis revealed that RC organoids exhibit a higher expression of OXPHOS-related genes compared to glycolysis-related genes, suggesting a metabolic shift towards oxidative phosphorylation. To further characterize these metabolic states, we developed a novel protocol, which enables precise and reliable quantification of mitochondrial oxygen consumption (OCR). Our findings emphasize the importance of controlling the physical culture environment to ensure consistent outcomes in human neurodevelopmental models while showcasing the impact of our newly established culturing strategy on implementing state-of-the-art technologies.

## RESULTS

### Use of near-microgravity conditions for the generation of brain organoids

To minimize the mechanical stress during key stages of differentiation, we made use of a Clinoreactor system (https://celvivo.com), herein called Rotating Chamber (RC), which enables precise control over fluid dynamics during organoid development. Unlike traditional bioreactors or spinning platforms that expose the developing tissues to fFSS, the RC utilizes gentle rotational dynamics to achieve near-microgravity conditions, reducing shear stress while maintaining optimal nutrient and oxygen exchange (Wrzesinski & Fey, 2018) (**Fig 1A**). Computational fluid dynamics (CFD) analysis performed for the RC demonstrated an even distribution of shear forces within the culture chamber (**Fig 1B**). Shear forces remained consistently low (< 14 mPa), a range suitable for promoting organoid development without causing mechanical disruption (Saglam-Metiner *et al*., 2023; Velasco *et al*, 2020). Additionally, velocity and shear stress analyses confirmed stable flow dynamics across constructs with different densities (**Fig 1C, movie EV1** and **Methods**). The shear stress levels remained low across all tested conditions, confirming the RC’s ability to maintain a reproducible and mechanically stable culture environment, suitable for generating reproducible brain organoids under optimized biomechanical conditions (**Fig 1D**).

**Fig 1.**
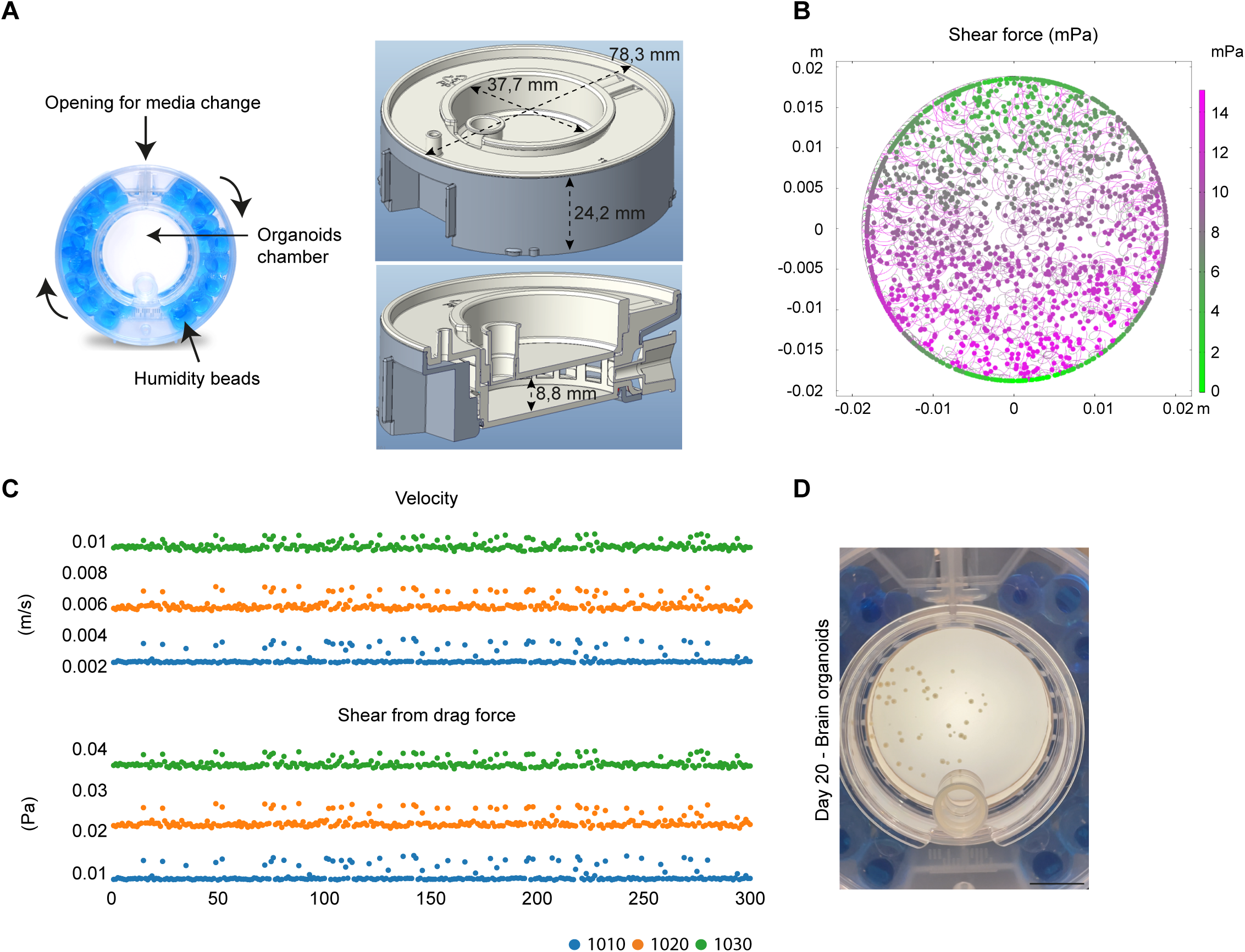
Regulation of shear force affects organoids development. **(A)** Schematic representation of the Rotating Chamber (RC), illustrating its rotational dynamics designed to minimize fluid flow shear stress (fFSS) while ensuring uniform nutrient and oxygen distribution. The representation highlights key features, including the opening for media exchange, the organoid culture chamber, and the surrounding humidity beads to maintain optimal culture conditions. The right panels provide detailed dimensions of the CR apparatus, with an outer diameter of 78.3 mm and a height of 24.2 mm. The inner culture chamber has a diameter of 37.7 mm and a height of 8.8 mm, providing ample space for organoid growth. **(B)** Computational Fluid Dynamics (CFD) (COMSOL Multiphysics 6.2) simulation depicting the distribution of shear forces within the culture chamber. The shear force map reveals consistently low values (<20 mPa), indicating a mechanically stable environment conducive to organoid development. **(C)** Velocity and shear stress analysis for constructs with different density (Generated in COMSOL Multiphysics 6.2 by Resolvent Denmark PS, Maaloev, Denmark; relative mass 1010, 1020, and 1030 where 1000 is density/relative mass of surrounding medium). The top graph shows the velocity distribution, while the bottom graph illustrates shear stress from drag force. Both analyses confirm the stability and reproducibility of the CR system in maintaining low-shear conditions for all individual objects within organoid chamber (objects 0 to 300). **(D)** Representative brightfield image of brain organoids cultured in the RC system at day 20. Scale bar: 1 mm.

### Differentiation of brain organoids in RC conditions

To generate brain organoids, we adapted a previously published protocol for dorsal forebrain differentiation (Khan *et al*, 2020; Sloan *et al*, 2018) (**Fig 2A**, top). This protocol was chosen as it is matrix-free, and therefore aligns with our objective to avoid the variability and presence of undefined factors associated with ECM-based products (Jain *et al*., 2023; Long *et al*., 2018; Martins-Costa *et al*., 2023).

**Fig 2.**
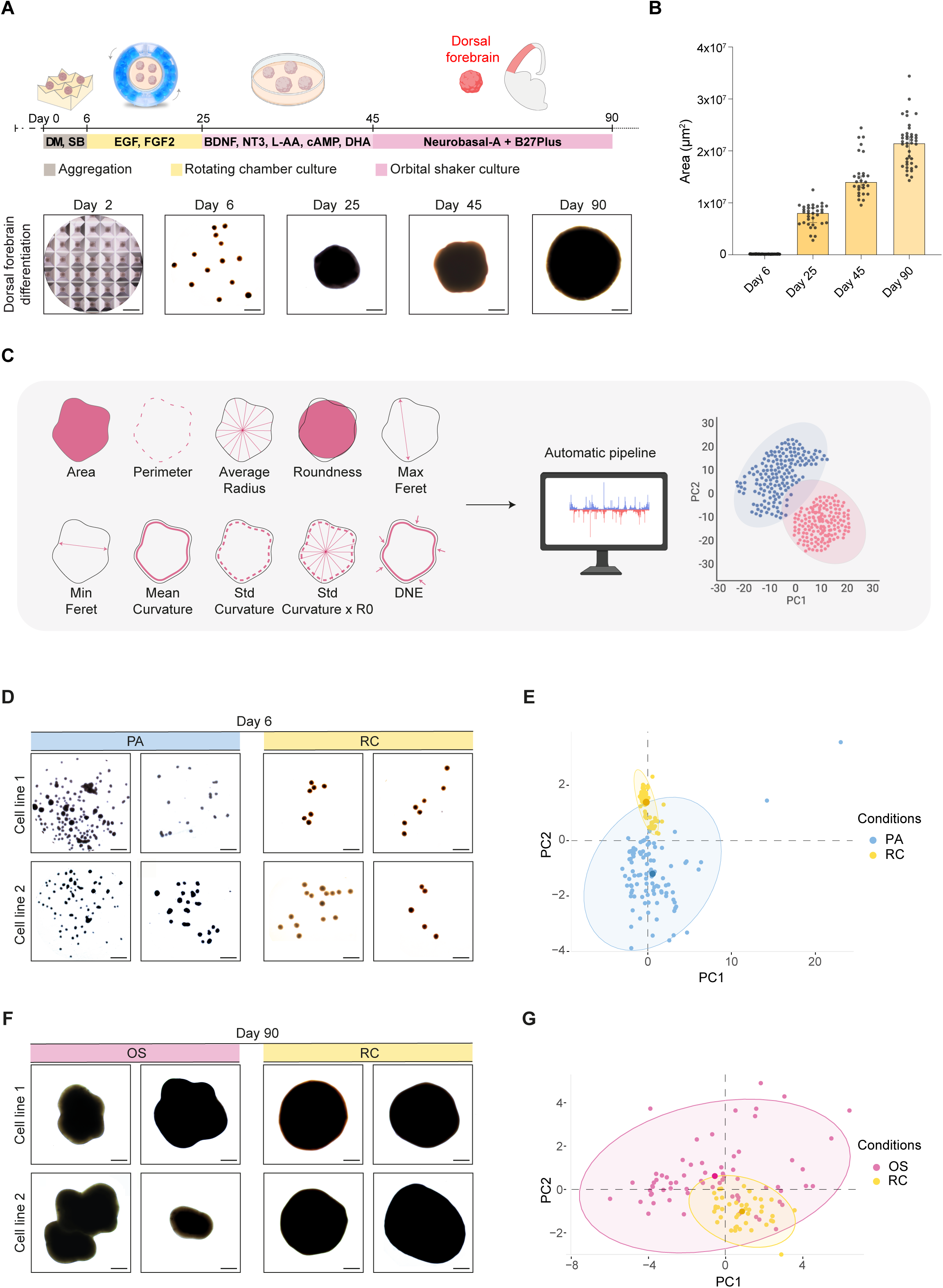
RC organoids exhibit reduced morphological variability across batches and cell lines. **(A)** Schematic representation of the experimental timeline for RC organoid generation, from aggregation (Day 0) to maturation (Day 90). Representative brightfield images illustrate the morphological progression of dorsal forebrain organoids at different time points. Scale bars: 500 µm. **(B)** Quantification of organoid areas at different time points, showing a significant increase in size from Day 6 to Day 90. Data are presented as mean ± SEM. **(C)** Overview of the automated morphological analysis pipeline. Ten morphological parameters, including area, perimeter, average radius, roundness, Feret diameters, curvature metrics, and Dirichlet normal energy (DNE), were extracted from organoid images. The dataset was processed using an automated Python-based code, and principal component analysis (PCA) was applied to assess variability across conditions. **(D)** Representative brightfield images of embryoid bodies (EBs) at Day 6, comparing PA and RC protocols across two independent iPSC lines. Each picture represents EBs of different batches. PA EBs display a heterogeneous morphology with frequent aggregation, while RC EBs exhibit a more uniform, spherical shape. Scale bars: 500 µm. **(E)** PCA analysis of morphological parameters at Day 6, comparing PA (blue) (n = 102, five batches) and RC (yellow) (n = 88, three batches) conditions. The confidence ellipses englobe 95% of the EBs within the same condition and its center, indicating the mean of all the data, is highlighted with a darker colour. **(F)** Representative brightfield images of organoids at Day 90, comparing OS and RC protocols across two iPSC lines. Each picture represents organoids from different batches. OS organoids exhibit irregular and lobulated morphologies, whereas RC organoids maintain a more uniform, spherical shape. Scale bars: 500 µm. **(G)** PCA analysis of morphological parameters at Day 90, comparing OS (pink) (n = 67, five batches) and RC (yellow) (n = 41, five batches) conditions. Each dot represents one organoid. The confidence ellipses englobe 95% of the organoids within the same condition and its centre, indicating the mean of all the data, is highlighted with a darker colour.

First, we generated embryoid bodies (EBs) from hiPSCs and extended the initial aggregation phase to six days, with minimal disturbance, in ultra-low attachment microwell culture plates (AggreWell™) to reduce the risk of early fusion events. The EBs were then transferred into the RC apparatus for the entire neural induction phase (Days 6–25) in the presence of EGF and FGF2. Within the RC system, the organoids remained gently suspended in the chamber, and exhibited a smooth, uniform, and spherical morphology (**Fig 2A** and **movie EV2)**.

For the differentiation phase (Days 25-45), organoids were transferred to low attachment tissue culture plates maintained on an orbital shaker, in the presence of brain-derived neurotrophic factor (BDNF), neurotrophin-3 (NT3), cyclic AMP (cAMP), L-ascorbic acid (L-AA), and docosahexaenoic acid (DHA). Thereafter (Days 45-90), organoids were kept in maturation medium (Neurobasal-A, B27 Plus). Morphologically, organoids transitioned from compact EBs at day 2 to more organized and enlarged structures by day 6 (**Fig 2A** bottom). As differentiation progressed, concomitant with a steady growth in size, the organoids maintained a consistent spherical shape, which is critical for uniform development of the different organoids within the same batch, consistent with ongoing proliferation and maturation (**Fig 2A** bottom). Organoid area increased significantly over time, with the largest values observed at day 90 (**Fig 2B**).

### RC organoids exhibit greater morphological consistency across different batches and cell lines

Morphological parameters are critical indicators of organoid health, quality, reproducibility and functionality (Chiaradia *et al*., 2023; Lancaster *et al*., 2017; Øhlenschlæger *et al*., 2023). We developed an automated Python-based pipeline to extract and quantify 10 distinct, previously established morphological parameters (Chiaradia *et al*., 2023) (**Fig 2C** and **Methods**). This approach enables objective, scalable, and reproducible morphometric analysis, minimizing user bias and ensuring consistency across experiments.

We conducted this analysis at two different stages of organoid development, days 6 and 90, comparing two alternative culture protocols across two different hiPSC lines. In the first comparison, we employed the classic method of organoid culture (Sloan *et al*., 2018), referred to as PA protocol, and our RC protocol. The PA protocol involves the aggregation in ultra-low-attachment plates for the first day, followed by 6 days of dual-SMAD inhibition (Khan *et al*., 2020; Sloan *et al*., 2018) (**Fig EV1A**). This method involves the simultaneous inhibition of two key signaling pathways: the BMP (Bone Morphogenetic Protein) and the TGF-β (Transforming Growth Factor Beta) pathways, both involving SMAD proteins as intracellular signaling mediators. Conversely, the RC protocol consists of 6 days of aggregation in AggreWell^TM^ plates, using the same media formulation (**Fig 2A** and **Fig EV1A**).

At day 6, brightfield images revealed considerable differences in the morphology of PA and RC EBs (**Fig 2D**). EBs grown with the PA protocol tend to fuse, resulting in cellular aggregates of various sizes and shapes. By contrast, EBs grown with RC protocol exhibit a spherical morphology, a more uniform size, and rarely fuse together. Principal component analysis (PCA) of all the morphometric measurements shows that PA EBs exhibit a larger confidence ellipse than RC EBs (**Fig 2E**), indicating greater morphological heterogeneity. The contribution of each morphological parameter, calculated as the cosine² across the principal components (PCs) of the PCA, is summarized in a correlation plot (**Fig EV1B**). Each parameter contributes strongly to the first two dimensions of the PCA, except for roundness, which is represented to a lesser extent, and the mean curvature, which is better characterized by the PC3. Except for perimeter, all other morphological parameters are significantly different between the two protocols (**Fig EV1C**). Overall, RC EBs are larger than PA EBs, with statistically significant higher median areas and average radii. Similar observations are noted for the minimum and maximum Feret diameters. Importantly, the median roundness values for PA and RC EBs are 85.5% and 96.9%, respectively, indicating that the RC protocol generates more spherical EBs (**Fig EV1C** and **Dataset EV1**). In addition, PA EBs display a significantly wider distribution across all morphological variables (**Dataset EV2**).

To assess the inter-batch variability, we quantified the absolute value of the relative median absolute deviation (MAD) (**Fig EV1D** and **Dataset EV3**). Except for the standard curvature and the Dirichlet normal energy (DNE), all absolute relative MADs indicate a significant difference in variability across batches, with the PA protocol displaying a greater value of MAD (Mann-Whitney test, p < 0.05). Taken together, these results suggest that the RC EBs demonstrate higher inter-batch morphological reproducibility compared to the PA EBs.

To further investigate the reproducibility of morphological features across different cell lines, we analysed the inter-line variability within the RC and PA protocols. RC EBs exhibit a more consistent distribution of morphological measurements across both lines, with no significant differences in area, perimeter, or average radius (**Fig EV1E)**. By contrast, PA EBs display greater differences between the two lines, particularly in roundness and curvature-related parameters. The median roundness values show a more pronounced difference between the two lines in the PA protocol compared to the RC protocol, indicating that the RC system facilitates greater morphological uniformity across cell lines with different genetic backgrounds.

The inter-line variability was further quantified through standard curvature and DNE measurements, revealing that RC EBs exhibit slightly higher fluctuations **(Fig EV1E)**. This indicates that while the RC protocol enhances inter-batch reproducibility, subtle morphological differences across cell lines may persist due to inherent genetic variability. The results suggest that the RC protocol not only enhances inter-batch reproducibility but also mitigates morphological variability across different cell lines. Importantly, despite minor genetic differences, RC EBs maintain a more stable morphology across lines compared to PA EBs, demonstrating the robustness of the RC protocol in mitigating morphological heterogeneity during early differentiation.

Given the results obtained with the 6 days aggregation protocol, we performed a second comparison between the RC protocol and a more traditional approach for organoid neural induction, which utilizes orbital shaker, referred to as OS protocol. This protocol is identical to the RC protocol, except for the different culture vessels used during the neural induction phase from day 6 to day 25 (**Fig EV2A**). Organoids were generated from two different hiPSC lines and harvested for analysis at a more mature stage (day 90). OS organoids from both lines at day 90 exhibit irregular and multilobate morphology.

Conversely, RC organoids display consistently more regular and spherical shapes (**Fig 2F**). Considering all morphometric parameters in the PCA analysis, OS organoids present a broader ellipse compared to RC organoids (**Fig 2G**), indicating greater morphological heterogeneity. The contribution of each morphological parameter reveals that the features are well represented by the first two dimensions of the corresponding PCA plot, except for roundness and standard curvature (**Fig EV2C**). This is further illustrated in the bar plots of each variable (**Fig EV2C**). Notably, the median roundness values of OS and RC organoids are 78.0% and 90.2%, respectively. Among the measured morphological parameters, area, average radius, minimum and maximum Feret diameters significantly differ between the two groups, indicating that the RC protocol tends to generate larger organoids. Furthermore, the OS protocol produces organoids with a wider distribution of their morphometric values compared to RC organoids for all features, except for roundness and standard curvature (**Dataset EV2**). The inter-batch variability is illustrated with the absolute relative MAD (**Fig EV2D** and **Dataset EV4**). OS batches show greater variability in perimeter and roundness values compared to RC batches. These findings suggest that the RC protocol allows the generation of morphologically more reproducible organoids, even during long-term culture, by standardizing morphological parameters and reducing inter-batch variability, across different cell lines and advanced stages of differentiation.

To further assess the positive effect of controlling the shear force and fluid dynamics on organoid morphology at later stages of differentiation, we analysed the inter-line variability at day 90. The comparison was performed between RC and OS organoids generated with two protocols across two independent hiPSC lines. OS organoids display higher morphological variability between the two lines, particularly in area, perimeter, average radius and curvature-related features, whereas RC organoids exhibit a more uniform profile. Notably, roundness values remain significantly higher in RC organoids across both lines, reinforcing the ability of the RC system to maintain a more consistent structural organization (**Fig EV2E**). Although some differences between the two hiPSC lines persist, the RC protocol significantly mitigates morphological variability compared to OS, highlighting its robustness in generating reproducible organoids.

These findings indicate that the RC methodology effectively standardizes morphometric parameters while also reducing inter-line variability at later differentiation stages. By integrating automated morphological analysis with PCA and direct statistical comparisons, we demonstrate that the RC protocol ensures consistent organoid quality and reproducibility, establishing a reliable framework for downstream applications.

### RC organoids display dorsal forebrain identity

To assess the differentiation and cellular identity in organoids generated using the RC protocol, we harvested day 20 organoids (**Fig 3A**) and subjected them to immunohistology analysis **(Fig 3B).**

**Fig 3.**
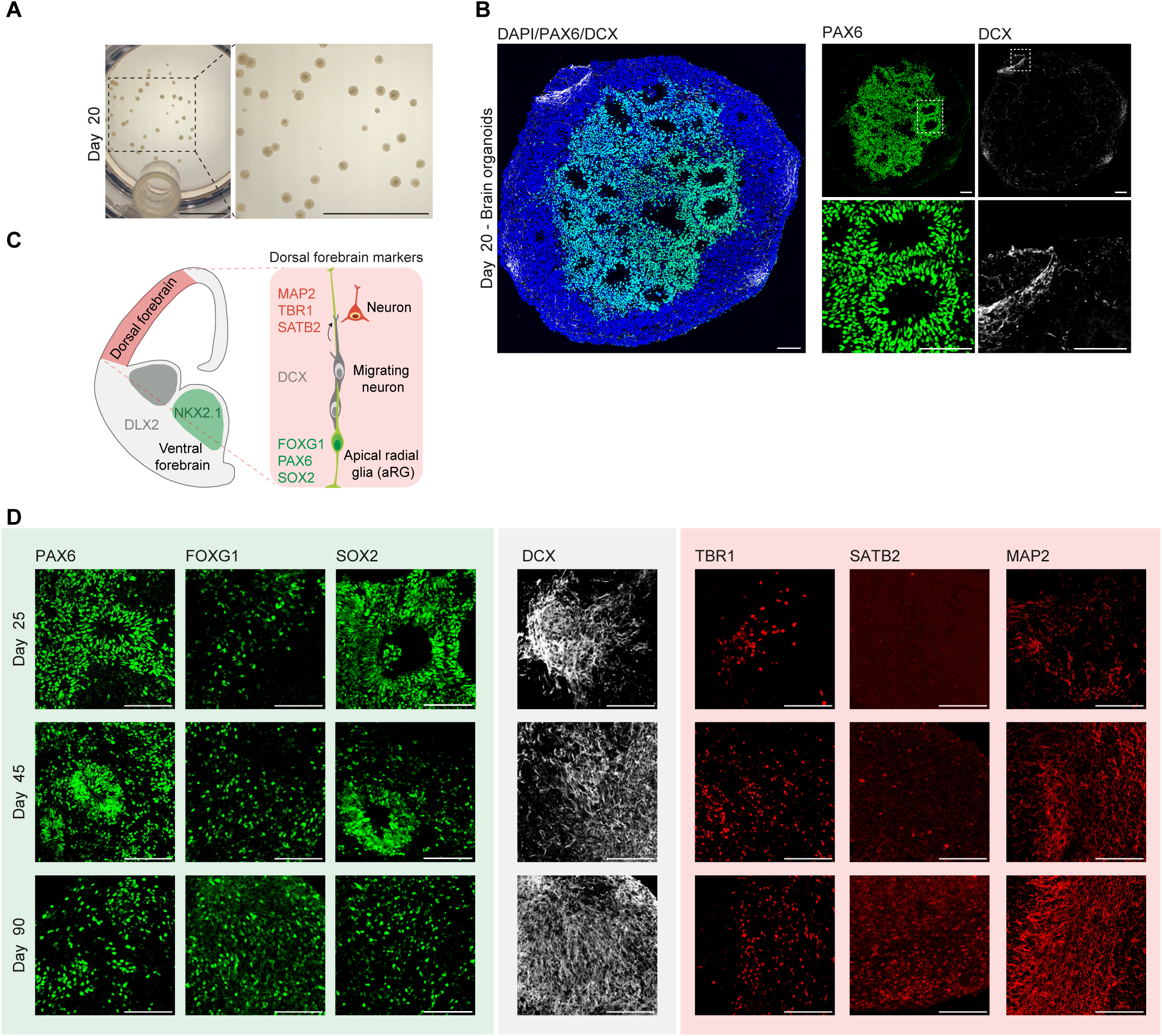
RC organoids exhibit dorsal forebrain identity. **(A)** Brightfield images of day 20 organoids cultured in the RC system, showing uniform morphology and size distribution. Scale bars: 1 mm. **(B)** Immunofluorescence images of day 20 organoids, stained for PAX6 (green), DCX (white), and DAPI (blue). The main panel provides an overview of the organoid structure, while high-magnification insets, highlighted by a dashed-line rectangle, show the distribution of PAX6+ progenitors and DCX+ migrating neurons. Scale bars: 100 μm. **(C)** Schematic representation of dorsal and ventral forebrain markers, illustrating marker expression patterns in organoids. **(D)** Immunofluorescence staining at days 25, 45, and 90, showing expression of PAX6, FOXG1, and SOX2 (green), DCX (white), and TBR1, SATB2, and MAP2 (red) across different time points. Scale bars: 100 μm.

This analysis revealed the presence of numerous neuroepithelial rosettes at the innermost parts of the organoids. These cellular organizations are a hallmark of early neural patterning and are predominantly positive for PAX6, which is a marker of dorsal forebrain progenitors. At the periphery and throughout the organoids, we identified foci of DCX-positive cells, which is a marker for newly differentiated immature neurons (Braun *et al*, 2023) (**Fig 3B** right). This confirmed that the RC system preserves both the external morphology and the inner cytoarchitecture of organoids, ensuring uniform tissue organization.

Next, we profiled organoids at day 25, 45 and 90 to follow the temporal evolution of the tissue patterning using a panel of marker proteins specifically expressed during dorsal forebrain development (**Fig 3C**). These proteins include SOX2, a marker for multipotent neural stem cells; FOXG1, a transcription factor critical for forebrain specification; PAX6, a marker of radial glia and neural progenitors; SATB2, a marker of upper-layer cortical neurons; and TBR1, a marker of deep-layer cortical neurons. To evaluate neuronal maturation, we detected DCX and MAP2. Additionally, we assessed the expression of proteins expressed in ventral forebrain, including NKX2.1 and DLX2, which are associated with medial ganglionic eminence (MGE) identity and interneuron progenitors respectively and therefore should not be present (Braun *et al*., 2023) (**Fig 3D** and **Fig EV3**).

At day 25, FOXG1, PAX6, and SOX2 were prominently expressed, indicating neural progenitor populations with dorsal forebrain identity **(Fig 3D** top left). DCX expression also began to emerge, demonstrating early neuronal specification (**Fig 3D** top middle) and expression of TBR1, a marker of early born deep layer neurons was also detected (**Fig 3D** top right). At day 45, TBR1 and MAP2 expression increased, reflecting the ongoing maturation of cortical neurons, and SATB2 positive cells corresponding to later-born cortical neurons start to be detected (**Fig 3D** middle right). PAX6 and SOX2 expression persisted at ‘rosette-like’ progenitor regions (**Fig 3D** middle left). By day 90, the expression of neuronal markers MAP2 and DCX dominated, with cortical neuron markers TBR1 and SATB2 reaching the highest abundance compared to the earlier time points. Notably, FOXG1 expression was maintained throughout differentiation, further confirming forebrain identity. Importantly, ventral forebrain markers NKX2.1 and DLX2 were undetectable across all the timepoints (**Fig. EV3A**), underscoring the regional specificity of RC-harvested organoids to the dorsal forebrain. The absence of ventral markers, combined with the expression of dorsal markers, highlights the robustness of our protocol in driving region-specific organoid differentiation.

### RC organoids recapitulate consistently cell types of the developing human brain

To more comprehensively characterize the identity of the RC organoids, we evaluated their cellular composition at day 90 using single-cell RNA sequencing technology (**Fig 4A**). This time point was chosen as it represents a late stage of organoid maturation, where cellular diversity is well-established, allowing for a comprehensive evaluation of neuronal differentiation and progenitor maintenance (Khan *et al*., 2020; Velasco *et al*., 2019).

**Fig 4.**
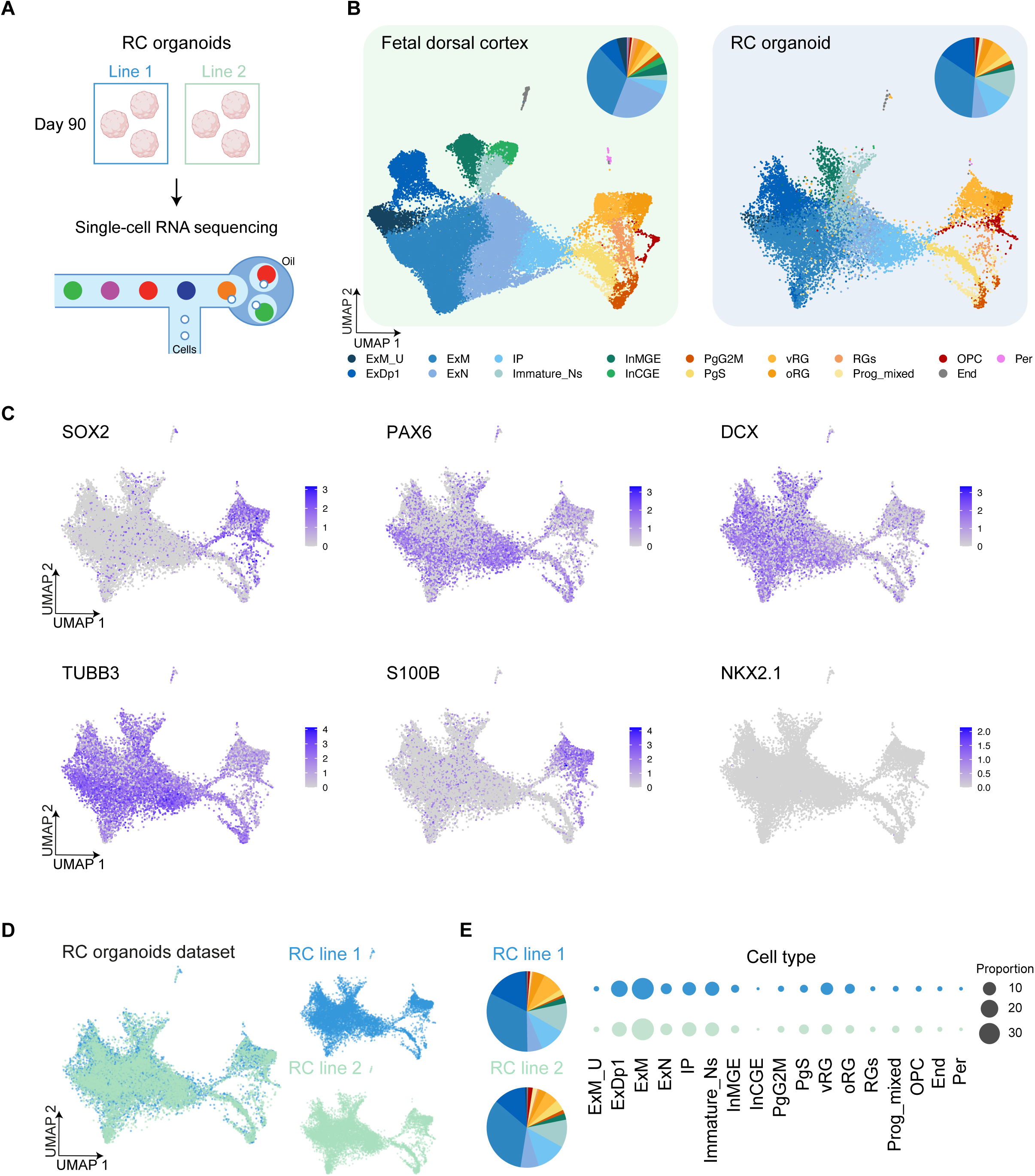
RC organoids recapitulate cortical cell types and exhibit inter-line cellular reproducibility. **(A)** Schematic representation of the experimental workflow. RC organoids from two independent iPSC lines were harvested at day 90, dissociated into single cells, and subjected to single-cell RNA sequencing. **(B)** UMAP representation of the fetal dorsal cortex (left) (Polioudakis *et al*., 2019) and RC organoids (right), illustrating the clustering of cell populations. The pie charts indicate the proportional distribution of different cell types in each dataset. **(C)** Feature plots displaying the expression patterns of key neural progenitor and neuronal markers, including SOX2 (neural progenitors), PAX6 (dorsal progenitors), DCX (migrating neuroblasts), TUBB3 (neuronal marker), S100B (radial glia and oligodendrocyte progenitors), and NKX2.1 (ventral forebrain marker). **(D)** UMAP representation of the RC organoid dataset, separated by iPSC lines (RC line 1 and RC line 2), demonstrating consistency between replicates. **(E)** Proportional representation of cell types across the two RC organoid lines, visualized through dot plots and pie charts, confirming the reproducibility of the differentiation process. Cell types include: ExM_U (maturing excitatory upper enriched), ExDp1 (excitatory deep layer 1), ExM (maturing excitatory), ExN (migrating excitatory), IP (intermediate progenitors), Immature_Ns (immature neurons), InMGE (interneuron medial ganglionic eminence), InCGE (interneuron caudal ganglionic eminence), PgG2M (cycling progenitors in G2/M phase), PgS (cycling progenitors in S phase), vRG (ventricular radial glia), oRG (outer radial glia), RGs (radial glial populations), Prog_mixed (mixed progenitors), OPC (oligodendrocyte progenitor cells), End (Endothelial cells) and Per (pericytes).

As part of the preprocessing step, we applied the computational algorithm *Gruffi* (Vertesy *et al*, 2022) (see **Methods**), which employs granular functional filtering to remove stressed cells in an unbiased manner. After filtering, a total of 17’738 high-quality cells were retained for further analysis, providing a refined dataset for downstream comparisons (**Fig EV4A**). We annotated the single-cell transcriptomic data from RC organoids using a reference human fetal brain dataset (Polioudakis *et al*, 2019) (see **Methods**). The fetal dataset includes distinct progenitor populations, including cycling progenitors in both G2/M (PgG2M) and S (PgS) phases, and mixed progenitors (Prog_mixed). Radial glial cells (RGs) include outer radial glia (oRG) and ventricular radial glia (vRG), while excitatory neuronal populations comprise maturing excitatory upper enriched (ExM_U), excitatory deep layer 1 (ExDp1), maturing excitatory (ExM), migrating excitatory (ExN), and immature neurons (Immature_Ns). In addition, intermediate progenitors (IP), early oligodendrocyte progenitor cells (OPCs), as well as interneuron populations from both the medial ganglionic eminence (InMGE) and caudal ganglionic eminence (InCGE), including a small population of endothelial (End) cells and pericytes (Per), were identified (**Fig. 4B** left). Our analysis revealed that RC organoids closely recapitulate all the major cell types of the mid-gestation human cortex, showing a highly similar cellular distribution between RC organoids and the fetal dorsal cortex (**Fig. 4B** right).

To further validate the cellular identity of these populations, we examined the expression of specific markers within the UMAP representation (**Fig 4C**). Neural progenitor populations exhibited strong expression of SOX2 and PAX6, key markers of dorsal forebrain progenitors. Neuronal populations displayed robust expression of DCX and TUBB3, confirming the presence of both migrating neuroblasts and maturing neurons. Additionally, S100B was expressed in radial glia and OPC. As expected, the ventral forebrain marker NKX2.1 was not expressed, reinforcing the regional specificity of RC organoids toward the dorsal forebrain identity.

To investigate the functional profiles of the cell types, present in the RC organoid dataset, we identified the most significant marker genes for each cell type (see **Methods**) and performed Gene Ontology (GO) enrichment analysis for each cell population (**Fig EV4B**). The analysis confirmed that the RC organoid cell types are associated with processes relevant to brain development. For example, ExM were enriched for terms such as “axonogenesis” and “regulation of synapse organization”. ExM_U and Immature_Ns were enriched for “forebrain development” while progenitor populations, such as oRG, showed enrichment for “regulation of neuron projection development”, while processes including “chromosome segregation” were prominent in proliferative populations. These results emphasize the functional alignment of RC organoid cell types with their *in vivo* counterparts (Polioudakis *et al*., 2019), further supporting the physiological relevance of RC-generated organoids.

We next evaluated the inter-line reproducibility by comparing the proportions of cell types between the two RC lines, using the fetal brain samples as reference (Polioudakis *et al*., 2019). The UMAP representation demonstrated a strong overlap across all cell types between the two hiPSC lines (**Fig 4D**). The proportions of cell types across the two different lines and gestational weeks are strongly overlapping (**Fig 4E**), consistently demonstrating that the cell type distributions were comparable between the two RC lines.

Together, these analyses confirm that the RC protocol generates highly reproducible cellular compositions across different genetic backgrounds, with the proportions of key progenitor and neuronal populations being remarkably consistent. Additionally, these findings highlight the inter-line reproducibility of cellular composition in RC organoids.

### Metabolic profiling of RC organoids

Energy metabolism has recently emerged as a key focus area in neuroscience, particularly in the context of neurodevelopmental disorders, where metabolic dysfunctions are increasingly implicated in a wide range of neurological conditions (Camandola & Mattson, 2017; Gandal *et al*, 2018; Kanellopoulos *et al*, 2020; Mariano *et al*, 2023). Moreover, metabolic regulation plays a crucial role in neurodevelopment, influencing processes such as progenitor proliferation, neuronal differentiation, and synaptic activity (Badal *et al*., 2019; Iwata *et al*., 2023; Iwata *et al*., 2020; Khacho & Slack, 2018). However, recent studies have highlighted metabolic heterogeneity in brain organoids, demonstrating how shifting between OXPHOS and glycolysis impacts differentiation outcomes (Øhlenschlæger *et al*., 2023). Given the transcriptional signatures linked to mitochondrial function and oxidative metabolism observed in RC organoids (**Fig EV4B**), we sought to investigate whether specific metabolic pathways, OXPHOS and glycolysis, were differentially regulated across cell types. The analysis showed that OXPHOS-related genes are broadly expressed across multiple cell types, with particularly high expression in progenitors (PgS, PgG2M) and radial glia (oRG, vRG, RGS), whereas glycolysis-related genes exhibit a more evenly distributed but lower expression overall (**Fig 5A**). This transcriptional evidence indicates that RC organoids predominantly rely on mitochondrial respiration for energy production. Despite the growing interest, measuring oxygen consumption rates (OCR), a critical indicator of mitochondrial activity and metabolic health, is not trivial in brain organoids and, to our knowledge, not well-established. To address this gap, we established a novel protocol using the Oroboros O2K system (Bird *et al*, 2019) to measure mitochondrial oxygen consumption in the whole RC organoids. We dissociated a pool of three organoids for each of the two lines at day 90 and proceeded with the OCR protocol (see **Methods**), which captures key metabolic states, including OXPHOS, electron transport system (ETS) capacity, ROX and COX activity across distinct mitochondrial complexes (**Fig 5B**). Example traces (**Fig 5B**, left) illustrate the profile of a well responding pool, providing detailed insight into the functional contributions of complexes I, II, and IV. Quantitative analysis (**Fig 5B**, right) revealed that RC organoids exhibit a significantly overall higher OCR compared to OS organoids, indicating a metabolic shift toward enhanced OXPHOS. Notably, OCR levels remained highly consistent across batches and cell lines, further supporting the robustness and reproducibility of mitochondrial function in RC organoids (**Fig 5B**, right).

**Fig 5.**
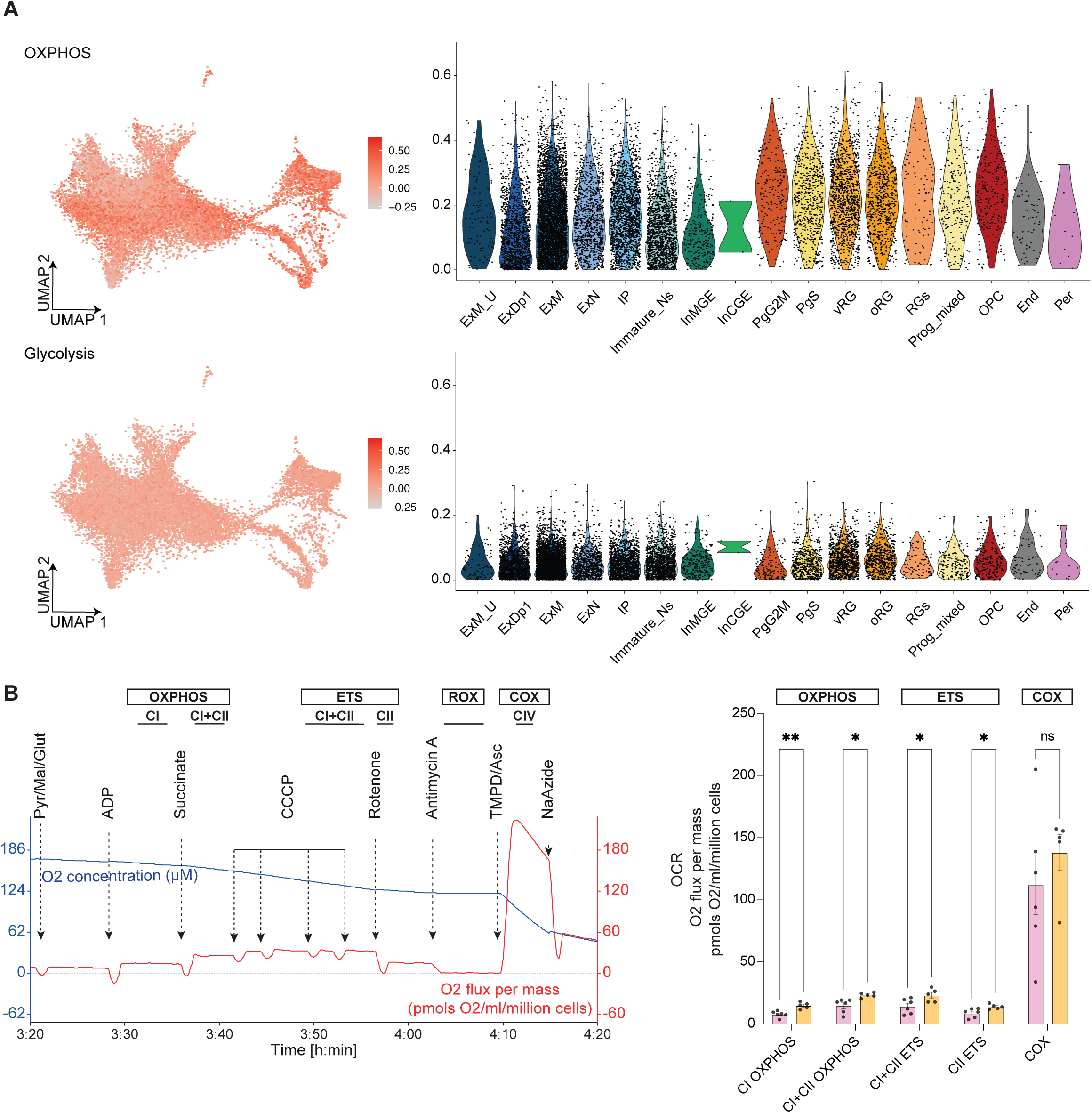
Metabolic profiling of RC organoids. **(A)** Single-cell transcriptomic analysis of oxidative phosphorylation (OXPHOS) and glycolysis-related gene expression across different cell types in RC organoids. UMAP plots on the left display the expression levels of OXPHOS and glycolysis genes, with intensity represented by a color scale. Violin plots on the right quantify the expression levels across different cell populations, including ExM_U (maturing excitatory upper enriched), ExDp1 (excitatory deep layer 1), ExM (maturing excitatory), ExN (migrating excitatory), IP (intermediate progenitors), Immature_Ns (immature neurons), InMGE (interneuron medial ganglionic eminence), InCGE (interneuron caudal ganglionic eminence), PgG2M (cycling progenitors in G2/M phase), PgS (cycling progenitors in S phase), vRG (ventricular radial glia), oRG (outer radial glia), RGs (radial glial populations), Prog_mixed (mixed progenitors), OPC (oligodendrocyte progenitor cells), End (Endothelial cells) and Per (pericytes). **(B)** Oxygen consumption rate (OCR) measurements in RC organoids at day 90 using high-resolution respirometry with the Oroboros O2K system. The panel on the left displays representative OCR traces, illustrating different mitochondrial respiration states: complex I oxidative phosphorylation (CI OXPHOS), combined complex I and complex II oxidative phosphorylation (CI + CII OXPHOS), combined complex I and complex II electron transport system (CI + CII ETS), complex II electron transport system (CII ETS), and cytochrome c oxidase activity (COX). Sequential compound additions, including ADP, succinate, CCCP, rotenone, antimycin A, and sodium azide, allow for the functional dissection of mitochondrial respiratory chain components. The panel on the right presents a quantitative comparison of OCR levels between RC organoids (yellow) and OS organoids (pink) normalized to cell number. Each dot represents a pool of three organoids from different batch. Statistical analysis was performed using multiple unpaired t-tests with Welch’s correction. Error bars represent the standard error of the mean (SEM). ns = not significant, * p < 0.05, ** p < 0.01, *** p < 0.001, **** p < 0.0001.

## DISCUSSION

Approximately 90% of clinical candidate drugs fail to progress from preclinical research to market approval (Seyhan, 2019). This high rate highlights a significant gap between preclinical models and real-world patient responses to treatments. Unlike conventional 2D cell cultures or rodent models, organoids more accurately mimic human physiology, offering a more reliable and scalable platform for assessing drug safety and efficacy (Antonica *et al*., 2022; Camp *et al*., 2015; Eichmuller & Knoblich, 2022; Kelley & Pasca, 2022; Kim *et al*., 2020; Lancaster *et al*., 2013; Luo *et al*., 2016; Pellegrini *et al*., 2020; Qian *et al*., 2019). However, the reproducibility of brain organoids remains a fundamental challenge in the field of neurodevelopmental modelling. Despite their potential to recapitulate critical features of human brain development, variability in morphological parameters, cellular composition, and differentiation outcomes often undermines their utility. Morphological parameters are critical indicators of organoid quality and reproducibility. Organoid cytoarchitecture not only reflects developmental health, but also influences downstream differentiation, regional patterning and functional properties (Chiaradia *et al*., 2023; Jain *et al*., 2023; Lancaster *et al*., 2017; Martins-Costa *et al*., 2023). Significant attention has been given to the biomechanical forces that act during the development of organoids. Indeed, these are essential drivers of tissue morphogenesis, influencing cell behaviour, spatial organization and differentiation.

The fFSS, a biomechanical force generated by media flow in dynamic culture systems, has long been recognized as a key factor influencing morphogenesis (Dahl-Jensen & Grapin-Botton, 2017; Goto-Silva *et al*., 2019; Hinton *et al*., 2022). While dynamic systems such as orbital shakers and bioreactors enhance nutrient and oxygen delivery, they also expose developing organoids to potentially disruptive fFSS. Excessive fFSS during critical phases of development can impair tissue integrity, disrupt cellular organization, and alter differentiation trajectories (Dardik *et al*., 2005; Gareau *et al*., 2014; Ismadi *et al*., 2014; Saglam-Metiner *et al*., 2023; Suong *et al*., 2021). In this context, multiple approaches have been explored to mitigate the effects of fFSS. These include optimizing spinning speed in bioreactors, using modified orbital shakers with reduced turbulence, or employing static culture systems that eliminate shear forces entirely. However, these approaches often come with drawbacks, such as uneven nutrient distribution in static cultures or residual variability in dynamic systems (Dardik *et al*., 2005; Gareau *et al*., 2014; Ismadi *et al*., 2014; Saglam-Metiner *et al*., 2023; Suong *et al*., 2021). Our study builds on this foundation by implementing the RC system, which drastically reduces fFSS thanks to gentle rotational dynamics.

This innovation creates a low-shear environment conducive to optimal 3D organoid development, achieving a better balance between nutrient delivery and mechanical stability. We demonstrated that morphological consistency ensures that organoids follow predictable developmental trajectories, reliably recapitulating spatiotemporal patterns of neurodevelopment. The RC system’s ability to maintain uniform spherical morphology, reduce early fusion events, and ensure even nutrient distribution highlights its efficacy in standardizing morphological parameters. Moreover, using the RC system, we achieved high inter-batch and inter-line reproducibility, as evidenced by consistent morphological parameters, cellular compositions, and metabolic profiles across independent experiments. Indeed, our novel OCR protocol demonstrated robust mitochondrial function with low variability, even across organoids derived from different cell lines. Importantly, our study reveals that organoids obtained with RC system using a guided protocol predominantly rely on OXPHOS rather than glycolysis, differently from what was described for organoids obtained using the standard guided protocol (Øhlenschlæger *et al*., 2023). This further corroborates the notion that biomechanical forces during key morphogenesis phases might have an impact on metabolic states, and consequently tissue fate.

This level of consistency enhances the utility of RC organoids for applications requiring reliable and reproducible models, such as high-throughput drug screening, disease modelling, and therapeutic testing. In addition, these findings emphasize the interplay between forces and morphology in organoid development and suggest that controlling biomechanical conditions can significantly improve reproducibility.

While our study highlights the advantages of the RC system for dorsal forebrain organoid generation, there are limitations that provide avenues for future improvements. Although the RC system effectively modulates biomechanical forces to enhance reproducibility, further optimization of the protocol is needed to achieve more precise and reliable patterning that closely mirrors human brain development. Emerging studies suggest that fine-tuning the molecular and environmental cues during differentiation is crucial for achieving more accurate regional identities. For instance, a recent study demonstrated that a dual SMAD and WNT inhibition pulse is both necessary and sufficient to establish robust and rich cortical cell repertoire that enables mirroring of fundamental molecular and cyto-architectural features of cortical development recapitulating the complexity and fidelity of cortical fate specification observed *in vivo* (Rosebrock *et al*, 2022). Incorporating such molecular adjustments into our protocol could further refine the developmental trajectories of organoids, ensuring more faithful recapitulation of spatiotemporal patterns and cell fate specification observed in the human brain. This underscores the importance of continuously integrating advances in biochemical modulation to complement biomechanical control.

In addition, while our OCR profiling demonstrated robust mitochondrial function across organoid batches, future experiments should aim to dissect in detail the metabolic heterogeneity across different neural populations. Specifically, investigating temporal shifts in energy metabolism and exploring how external cues, such as nutrient availability or disease-associated mutations, impact mitochondrial function will be crucial. Moreover, the metabolic states of the developing human brain remain largely unexplored, and further studies are needed to compare organoid-derived metabolic profiles with *in vivo* human data. Given the increasing recognition of metabolic dysfunctions in neurodevelopmental disorders, integrating metabolic profiling into standard organoid characterization will further enhance their utility as disease models.

In conclusion, by addressing the challenges of mechanical variability and morphological inconsistency, we have provided a robust framework for reproducible brain organoid generation, showing how the integration of biomechanical control, morphological quantification, and functional profiling offers a comprehensive approach for advancing brain organoid research. Our findings not only underscore the importance of reproducibility for downstream applications but also pave the way for future studies exploring the versatility of the RC system in generating region-specific organoids. This work positions the RC system as a valuable platform for studying neurodevelopment, modelling diseases, and testing therapeutic interventions.

## METHODS

### Human induced pluripotent stem cell (hiPSCs) culture

The human iPSC lines were kindly provided by Prof. Peng Jin (Emory University) and previously generated from skin biopsy samples of age-matched healthy males at Emory University (Kang *et al*, 2021). All iPSC lines were cultured in feeder-free conditions on Geltrex (Gibco)-coated cell culture dishes in E8 Basal Medium (Thermo Fisher Scientific) with 100 U/ml penicillin (Thermo Fisher Scientific) and 100 μg/ml streptomycin (Thermo Fisher Scientific), at 37 °C in 5% CO_2_. All human pluripotent stem cells lines were maintained below passage 50 and were negative for mycoplasma.

### Brain organoid generation and differentiation

The organoid generation protocol was adapted and modified for RC-harvested organoids from Sloan et al 2018 (Sloan *et al*., 2018) as follow. hIPSC at 70-80% of confluency were washed in DPBS-/-(Thermo Fisher Scientific) and incubated with Accutase (StemCell Technologies) at 37 °C for 5 minutes in a 5% CO_2_ incubator until the cells detached with gentle shaking. The cell suspension was transferred to a 15 mL conical tube containing an equal volume of DMEM/F12 (Thermo Fisher Scientific) to neutralize the enzymatic activity. Cells were centrifuged at 200 × g for 5 minutes, resuspended in the required volume of E8 medium (Thermo Fisher Scientific) supplemented with Rock Inhibitor (Y-27632) (Tebu-Bio), and adjusted to a final density of 2.5–3 million cells per mL. For EB formation, 1 mL of the single-cell suspension was transferred into a well of an AggreWell plate (StemCell Technologies), achieving a final volume of 1.5 mL per well. EBs were visible within 24 hours after seeding and were maintained in AggreWell plates until day 6. On day 1 post-seeding, half of the medium was replaced with Essential 6 medium supplemented with 2,5μM Dorsomorphin (Merck) and 10μM SB-431542 (R&D Systems). Half of the medium was changed daily until day 6, except for day 3. At day 6, EBs were transferred from the AggreWell plates to RC Rotating Chambers (CelVIVO) (previously prepared, as described below) containing 15 mL of Neurobasal A-Minus medium. This medium was composed of Neurobasal-A medium (Thermo Fisher Scientific) supplemented with B27 without vitamin A (2%) (Thermo Fisher Scientific), Glutamax (1%) (Thermo Fisher Scientific), EGF (20 ng/mL) (Thermo Fisher Scientific), FGF2 (20 ng/mL) (Thermo Fisher Scientific), and P/S. The medium was changed every two days and was maintained until day 25. From day 25 to day 45, organoids were transferred to orbital shakers in 6-or 10-cm dishes and cultured in Neurobasal A-Plus medium, consisting of Neurobasal-A medium supplemented with B-27 Plus (2%) (Thermo Fisher Scientific), Glutamax (1%), BDNF (20 ng/mL) (Thermo Fisher Scientific), NT-3 (20 ng/mL) (Thermo Fisher Scientific), L-ascorbic acid (200 μM) (Merck), DHA (10 μM) (Merck), and cAMP (50 μM) (Merck). The medium was changed three times per week. Starting from day 45, organoids were maintained in Neurobasal A-Plus medium supplemented with B-27 Plus (2%) and Glutamax (1%), without additional growth factors. The medium was changed twice a week to support long-term organoid survival and maturation.

### RC preparation

One day before transferring the EBs, RC systems (CelVIVO) blue beads were hydrated with 25 mL of sterile ultra-pure water, and the chambers were exposed to UV light overnight for sterilization. On the following day, the chambers were thoroughly washed with DPBS (-/-) and DMEM/F12 before adding 10 mL of Neurobasal A-Minus medium. The EBs were then carefully transferred into the prepared chambers and placed in the ClinoStar bioreactor for culture.

### PA and OS harvested organoids

Except for the following differences, the PA and OS protocols followed the same steps as those applied for RC organoids. In the PA protocol, EBs were generated after a one-day aggregation period and subsequently placed in standard cell culture dishes without rotational movement as previously described in Sloan et al and Khan et al (Khan *et al*., 2020; Sloan *et al*., 2018). In the OS protocol, EBs underwent a six-day aggregation period in AggreWells and, starting from day 6, were cultured on an orbital shaker to maintain dynamic conditions.

### Computational Fluid Dynamics (CFD) simulation

CFD simulations were performed using a Navier-Stokes-based solver to model fluid velocity and shear stress for constructs with different density (Generated in COMSOL Multiphysics 6.2 by Resolvent Denmark PS, Maaloev, Denmark; relative mass 1010, 1020, and 1030 where 1000 is density/relative mass of surrounding medium). Shear force distribution was computed based on the chamber’s geometry, and velocity profiles were extracted to evaluate the system’s stability. Data were plotted to compare velocity and shear stress from drag force across different conditions.

### Immunohistochemistry

Organoids were fixed in 4% paraformaldehyde for 1 hour at 4°C. After fixation, organoids were washed two times with PBS for 10 minutes each at room temperature and then incubated in 30% sucrose at 4°C until they fully sank. Organoids were embedded in a 7.5% gelatin (Sigma Aldrich) solution containing 10% sucrose (Sigma Aldrich) and sectioned at 20 μm using a Cryostat. For immunohistochemistry, sections were blocked and permeabilized in a solution of 0.3% Triton X-100 (Sigma Aldrich) with 3% goat serum (Sigma Aldrich) for 1 hour at room temperature. Primary antibodies were diluted in 0.3% Triton X-100 (Sigma Aldrich) with 3% goat serum (Sigma Aldrich) in PBS and incubated overnight at room temperature. Following primary antibody incubation, sections were washed three times in PBS. Secondary antibodies conjugated with Alexa Fluor 488, 568, or 647 (Invitrogen) were applied for 1 hour at room temperature, followed by three additional PBS washes. DAPI (Thermo Fisher Scientific) was included in the secondary antibody solution to stain nuclei. Finally, slides were mounted using Mowiol mounting medium.

The following antibodies were used to perform immunostainings: PAX6 antibody (901301; Biolegend; 1/500); FOXG1 antibody (ab18259; abcam; 1/200); SOX2 antibody (ab171380; abcam; 1/200); Doublecortin (E-6) antibody (sc-271390; Santa Cruz; 1/100); TBR1 antibody (ab31940; abcam; 1/200); SATB2 antibody (MA5-32788; Invitrogen; 1:100); MAP2 antibody (M4403; Sigma; 1/500); NKX2.1/TTF-1 antibody (ab133737; abcam; 1/200); DLX2 antibody (sc-393879; Santa Cruz; 1/200).

### Imaging

Images were acquired with Echo Revolve2-K2-1861 and Leica SP8 confocal microscopes and processed with Fiji ImageJ.

### Analysis of morphological parameters

Organoids were imaged at day 6 and day 90 using the Echo Revolve2-K2-1861 microscope (Bico Group) in brightfield mode with a 2x objective. The images were converted to TIFF format and analysed using a Python-based script. The organoid outlines, representing the Region of Interest (ROI), were automatically detected by the script and saved into new images for further verification. If the automatically detected contour did not perfectly match the organoid boundary, the threshold was adjusted within the script. In cases where a correct ROI could not be obtained, the organoid with abnormal segmentation was removed from the dataset by verifying its ID number in the result images. Across all conditions, the number of batches varied from three to five per time point and protocol, with a minimum of five organoids analysed per batch.

### Analysis related to morphological parameters

Ten morphological parameters, first described by Chiaradia et al (Chiaradia *et al*., 2023), were measured from brightfield images of brain organoids and are defined as the following:

Area (μm2): Area of the segmented organoid; Perimeter (μm): Perimeter of the segmented organoid; Average radius R0 (μm): Average distance of the contour from the center of the segmented organoid; Roundness (%): Calculated as 4π×*Area/Major axis*^2^. A roundness of 100% represents a perfect circle; Minimum Feret (μm): The minimum distance between any two points along the contour; Maximum Feret (μm): The maximum distance between any two points along the contour; Mean curvature (μm^-1^): Average of the curvature along the contour. The curvature is calculated as the inverse of the radius of the osculating circle and is computed as [(*dx ×dyy*) − (*dy ×dxx*)]/[(*dx^2^)+(dy*^2^)]^3/2^, where dx and dy are the first derivatives in x and y, and dxx and dyy are the second derivatives of the contour; Standard curvature (μm^-1^): Standard deviation of the curvature along the contour; Standard curvature x R0: Standard deviation of the curvature along the contour normalised by the average radius (R0); Dirichlet Normal Energy (DNE): Logarithm of the square of the variation of the normal n = (dy,-dx) of the contour projected on its tangent t = (dx, dy), where dx and dy are the first derivatives in x and y.

As the values did not follow a normal distribution and contained some extreme values (notably in PA and OS organoids, because of their inner heterogeneity), Mann-Whitney test (wilcox.test) was applied to compare the morphological parameters between two protocols. The measured values were represented with bar plots. The variability between each protocol was analysed with Brown-Forsythe test, using the LeveneTest function (center = median). The inter-batch variability was compared with Mann-Whitney test (wilcox.test) and was illustrated in bar plots with the Median Absolute Deviation (MAD), more precisely the absolute value of the MAD divided by the median (Dumitrascu *et al*, 2019). PCA graphs were performed using the PCA and fviz_par functions. To assess the importance of the contribution of each parameter according to the first three dimensions of the PCA plots, correlation plots were generated using the corrplot function on the cosine2 value (calculated with the get_pca_var function).

### Single-cell RNA sequencing

Organoids were cultured in Neurobasal A-Plus medium supplemented with B-27 Plus (2%) and Glutamax (1%) until harvesting at day 90. Pool of three organoids were separately incubated in Trypsin (Sigma Aldrich)/Accutase (StemCell Technologies) (1:1) containing 10 U/ml DNaseI (Thermo Fisher Scientific) in the gentleMACS™ Dissociator (Miltenyi Biotec) set at program NTDK1. After digestion the cell suspension was passed through a 70μm strainer. Samples were loaded to recover 16.000 cells onto a Chromium Next GEM Chip G Single Cell Kit (10x Genomics, PN-1000127) and processed through the Chromium controller to generate single-cell GEMs (Gel Beads in Emulsion). scRNA-seq libraries were prepared with the Chromium Next GEM Single Cell 3’ Kit v3.1 (10x Genomics, PN-1000269). Organoids libraries were pooled and sequenced using the AVITI 10X. A total of 166 million reads were requested per library to ensure sufficient coverage for downstream analysis.

### Single-cell RNA sequencing analysis

We aligned reads to GRCh38 human reference genome with Cell Ranger 7.2 (10x Genomics) using default parameters to produce the cell-by-gene, Unique Molecular Identifier (UMI) count matrix.

UMI counts were analysed using the Seurat R package v.5. Cells were filtered for a min. 1000 genes, maximal mitochondrial content of 10%. Resulting high quality cells were normalized (“LogNormalize”) for scaled for each cell to a total expression of 10K UMI. Doublets were removed using DoubletFinder, while unlabeled negative cells were retained as their distribution was similar to that of singlets. All analyses were conducted using doublet-free datasets.

### Stress cell identification and removal using *Gruffi*

To ensure data quality and remove stress-affected cells, we applied *Gruffi* algorithm (Vertesy *et al*., 2022). Stress-affected cells accounted for 15.1% of the dataset and were removed before further analysis. Following *Gruffi* filtering, the cleaned dataset contained 17,738 high-quality cells, ensuring that downstream analyses were performed on physiologically relevant populations.

### Integration with fetal brain and cell type annotation

To validate the identity of organoid-derived cell populations, we integrated our dataset with single-cell RNA-seq data from human fetal brains (Polioudakis *et al*., 2019). This allowed for a direct comparison between organoid and fetal cortical development, ensuring accurate cell type annotation.

### Visualization and analysis

UMAP projections were computed following Canonical Correlation Analysis (CCA)-based integration of organoid-derived single-cell RNA-seq data with human fetal brain datasets. Cluster identities were assigned using Seurat’s RenameIdents() function, and cell types were factorized to ensure consistent ordering across datasets. Feature plots were generated to visualize the expression of key marker genes, including SOX2, PAX6, NKX2-1, DCX, TUBB3, and S100B, across different cell types. Expression levels were mapped onto UMAP projections using Seurat’s FeaturePlot() function. To quantify cell-type proportions across batches and conditions, metadata was extracted and processed using group_by() and summarise() functions in tidyverse. Bar plots and stacked bar plots were created to visualize the relative frequencies of each cell type in different datasets. Additionally, pie charts were generated by plotting cell-type proportions in polar coordinates using ggplot2. Dot plots were used to compare cell-type proportions across experimental conditions, where dot size represents the percentage of cells in each category. The dataset was transformed using melt() and plotted using ggplot2. This combination of visualizations provided a detailed characterization of cell-type heterogeneity, lineage specification, and transcriptional states across organoid samples.

### GO enrichment analysis

To identify differentially expressed genes (DEGs) for each cell type within the dataset, we used the Seurat function FindAllMarkers() with the Wilcoxon test, setting a minimum percentage threshold of 0.2 and a log-fold change threshold of 0.25. This analysis identified marker genes specific to each cluster, which were then saved for further exploration.

For GO enrichment analysis, we performed Biological Process (BP) ontology enrichment using the clusterProfiler package. The marker genes were split by cluster and analyzed with the enrichGO() function, specifying the human genome annotation database (org.Hs.eg.db) as the reference. To account for background gene expression, the entire set of detected genes in the dataset was used as the universe for enrichment testing. The results were corrected for multiple comparisons using the Benjamini-Hochberg (BH) adjustment method, applying a q-value cutoff of 0.05. To refine the results and remove redundant GO terms, we applied the simplify() function with a similarity threshold of 0.7, selecting the most representative GO terms based on the lowest adjusted p-value. The results were then compiled into a summary data frame and visualized using ggplot2. The dot plot represents the most significantly enriched biological processes, with dot size reflecting the number of genes associated with each GO term and color intensity corresponding to the adjusted p-value.

### Metabolic module scoring analysis

To assess metabolic activity in organoid-derived cell populations, we computed module scores for oxidative phosphorylation (OXPHOS) and glycolysis using gene sets associated with their respective GO terms. The analysis was conducted in Seurat, applying the AddModuleScore() function to quantify pathway activation at the single-cell level. For OXPHOS, genes associated with GO:0006119 (oxidative phosphorylation) were retrieved from the org.Hs.eg.db database. Similarly, for glycolysis, genes linked to GO:0006096 (glycolytic process) were extracted. To avoid redundancy, only unique gene symbols were considered for scoring. The module scores were computed for each cell using AddModuleScore() in Seurat, which compares the expression of genes in the pathway against control gene sets. The resulting OXPHOS and glycolysis scores were visualized using UMAP feature plots, highlighting pathway activity across different cell types.

To further analyze metabolic heterogeneity, we generated violin plots comparing module scores across cell types, normalizing y-axis limits to ensure comparability between metabolic states. All computations were performed using the Seurat and ggplot2 packages, ensuring robust visualization and interpretation of metabolic trends in organoid datasets.

### Oxygen Consumption Rate (OCR) measurements via high-resolution respirometry

To assess mitochondrial function in RC OCR measurements were performed on 90-day-old organoids using a high-resolution oxygraph (Oroboros O2K). Organoids were maintained in Neurobasal A-Plus medium supplemented with B-27 Plus (2%) and Glutamax (1%) until harvesting. Pools of three organoids were separately dissociated into single-cell suspensions to ensure accurate normalization of OCR values per million cells. Organoids were enzymatically dissociated using a 1:1 mixture of Trypsin (Sigma Aldrich) and Accutase (StemCell Technologies), supplemented with 10 U/ml DNase I (Thermo Fisher Scientific). The suspension was incubated in the gentleMACS™ Dissociator (Miltenyi Biotec) under the NTDK1 program, followed by filtration through a 70 μm cell strainer to obtain a single-cell suspension. The resulting cells were resuspended in MiR05 buffer, a mitochondrial respiration-optimized medium, to maintain metabolic activity throughout the assay. The metabolic profiling protocol involved a series of sequential additions of specific compounds to dissect the activity of individual mitochondrial complexes and respiratory states. First, 10 μg/ml digitonin (Sigma Aldrich) was added to permeabilize the plasma membrane, ensuring access to substrates by the mitochondria. Then, 1 mM malate (Sigma Aldrich), 2.5 mM pyruvate (Sigma Aldrich), and 10 mM glutamate (Sigma Aldrich) were added to drive electron flow through complex I (CI) via the tricarboxylic acid (TCA) cycle. To stimulate oxidative phosphorylation (OXPHOS), 2.5 mM ADP (Calbiochem) was introduced, enabling ATP synthesis. Subsequently, 5 mM succinate (Sigma Aldrich) was added to activate complex II (CII) and assess combined CI + CII OXPHOS activity. To measure the maximal capacity of the electron transport system (ETS), 0.2 mM carbonyl cyanide m-chlorophenyl hydrazone (CCCP) (Sigma Aldrich), a proton uncoupler, was added stepwise until maximal respiration was achieved. To evaluate the contribution of CI and CII to respiration, 0.5 μM rotenone (Sigma Aldrich) and 0.5 μM antimycin A (Sigma Aldrich) were added, respectively. Finally, 2 mM sodium ascorbate (Sigma Aldrich) and 0.5 mM N,N,N′,N′-tetramethyl-p-phenylenediamine dihydrochloride (TMPD) (Sigma Aldrich) were used to stimulate cytochrome c oxidase (complex IV, COX), and 100 mM sodium azide (Sigma Aldrich) was added to inhibit COX and determine residual oxygen consumption (ROX). These measurements provided a comprehensive metabolic profile of the organoids, enabling the quantification of mitochondrial respiratory capacity, OXPHOS efficiency, and individual contributions of mitochondrial complexes to overall cellular respiration.

## STRUCTURED METHODS

| Reagent/Resource | Reference or Source | Identifier or Catalog Number |
| --- | --- | --- |
| **Media and Reagents**<br>**(cell culture)** |  |  |
| Essential 6<sup>TM</sup> Medium Kit | Thermo Fisher Scientific | Cat# A1516401 |
| Essential 8<sup>TM</sup> Medium Kit | Thermo Fisher Scientific | Cat# A1517001 |
| Neurobasal<sup>TM</sup> -A Medium | Thermo Fisher Scientific | Cat# 10888-022 |
| B-27<sup>TM</sup> Plus Supplement (50x) | Thermo Fisher Scientific | Cat# A3582801 |
| B-27<sup>TM</sup> Supplement minus vitamin A (50x) | Thermo Fisher Scientific | Cat# 12587010 |
| cAMP (N6,2'-O-dibutyryl adenosine-3',5' (cyclic)-monophosphate sodium salt) | Merck | Cat# D0627-25MG |
| DHA (cis-4,7,10,13,16,19-Docosahexaenoic acid) | Merck | Cat# D2534-25MG |
| DSM (dorsomorphin) | Merck | Cat# P5499-5MG |
| GlutaMAX<sup>TM</sup> Supplement 100X | Thermo Fisher Scientific | Cat# 35050038 |
| Human/Mouse/Rat BDNF Recombinant Protein, ReproTech<sup>®</sup> | Thermo Fisher Scientific | Cat# 45002 |
| Human EGF, Animal-free recombinant protein, ReproTech<sup>®</sup> | Thermo Fisher Scientific | Cat# AF10015 |
| Human FGF-basic (FGF-2/bFGF) (154 amino acid) Recombinant protein, ReproTech<sup>®</sup> | Thermo Fisher Scientific | Cat# 10018B |
| Human NT-3 Recombinant protein, ReproTech<sup>®</sup> | Thermo Fisher Scientific | Cat# 45003 |
| 2-phospho-L-ascorbic acid trisodium salt (L-AA) | Merck | Cat# 49752 |
| Penicillin-Streptomycin (10'000 U/ml) | Thermo Fisher Scientific | Cat# 15140122 |
| SB 431542 | R&D Systems | Cat# 1614 |
| Y-27632 | Tebu-Bio | Cat# S1049 |
| Accutase | StemCell technologies | Cat# 7922 |
| Anti-adherence rinsing solution | StemCell technologies | Cat# 07010 |
| DMEM/F-12, HEPES (1x) | Thermo Fisher Scientific | Cat# 11330032 |
| DPBS, calcium, magnesium (1x) | Thermo Fisher Scientific | Cat# 14040091 |
| DPBS, no calcium, no magnesium (1x) | Thermo Fisher Scientific | Cat# 14190144 |
| Geltrex™ LDEV-Free, hESC-qualifies,<br>Reduced growth factor basement<br>membrane matrix | Thermo Fisher Scientific | Cat# A1413301 |
| ReLeSR™ | StemCell technologies | Cat# 100-0483 |
| Trypsin, EDTA (0.25%), phenol red | Thermo Fisher Scientific | Cat# 25200056 |
| AggreWell™ 800 | StemCell technologies | Cat# 34815 |
| Clinoreactor (CelVivo plate) | CelVivo | Cat# 30003 |
| Ethanol | Merck | Cat# 51976 |
| Cellstar®, cell-repellent surface, 6 cm | Greiner Bio-One GmbH | Cat# 628979 |
| Cellstar®, cell-repellent surface, 10 cm | Greiner Bio-One GmbH | Cat# 664970 |
| **Antibodies** |  |  |
| PAX6 antibody (1/500) | Biolegend | Cat# 901301 |
| FOXG1 antibody (1/200) | Abcam | Cat# ab18259 |
| SOX2 antibody (1/200) | Abcam | Cat# ab171380 |
| Doublecortin (E-6)antibody (1/100) | Santa Cruz | Cat# sc-271390 |
| TBR1 antibody (1/200) | Abcam | Cat# ab31940 |
| SATB2 antibody (1/100) | Invitrogen | Cat# MA5-32788 |
| MAP2 antibody (1/500) | Sigma | Cat# M4403 |
| NKX2.1/TTF-1 antibody (1/200) | Abcam | Cat# ab133737 |
| DLX2 antibody (1/200) | Santa Cruz | Cat# sc-393879 |
| **Secondary antibodies** |  |  |
| Goat anti-Mouse IgG (H+L), Alexa<br>Fluor™ 488 | Invitrogen | Cat# A-11029 |
| Goat anti-Mouse IgG (H+L), Alexa<br>Fluor™ 546 | Invitrogen | Cat# A-11030 |
| Goat anti-Mouse IgG (H+L), Alexa<br>Fluor™ 647 | Invitrogen | Cat# A-21236 |
| Goat anti-Rabbit IgG (H+L), Alexa<br>Fluor™ 488 | Invitrogen | Cat# A-11034 |
| Goat anti-Rabbit IgG (H+L),<br>Alexa Fluor™ 546 | Invitrogen | Cat# A-11035 |
| Goat anti-Rabbit IgG (H+L),<br>Alexa Fluor™ 647 | Invitrogen | Cat# A-21244 |
| **Single-cell RNA sequencing** |  |  |
| Chromium Next GEM Chip G Single<br>Cell Kit | 10x Genomics | Cat# PN-1000127 |
| Chromium Next GEM Single Cell 3'<br>Kit v3.1 | 10x Genomics | Cat# PN-1000269 |
| **Oroboros O2K reagents** |  |  |
| MiR05 buffer | Oroboros Instruments | Cat# 60101-01 |
| Digitonin | Sigma Aldrich | Cat# D5628 |
| Malate | Sigma Aldrich | Cat# M1000 |
| Pyruvate | Sigma Aldrich | Cat# P2256 |
| Glutamate | Sigma Aldrich | Cat# G1626 |
| ADP | Calbiochem | Cat# 117105 |
| Succinate | Sigma Aldrich | Cat# S2378 |
| Carbonyl cyanide m-chlorophenyl<br>hydrazone (CCCP) | Sigma Aldrich | Cat# C2759 |
| Rotenone | Sigma Aldrich | Cat# R8875 |
| Antimycin A | Sigma Aldrich | Cat# A8674 |
| Sodium ascorbate | Sigma Aldrich | Cat# A4034 |
| N,N,N',N'-tetramethyl-p-phenylenediamine dihydrochloride<br>(TMPD) | Sigma Aldrich | Cat# T3134 |
| Sodium azide | Sigma Aldrich | Cat# S8032 |
| **Products** |  |  |
| Base Mould Disposable 15x15x06 mm | Kaltek | Cat# 2781 |
| DAPI | Thermo Fisher Scientific | Cat# 62247 |
| D-sucrose | Roth | Cat# 4621.1 |
| DPBS | Gibco | Cat# 14190-094 |
| Goat serum | Sigma Aldrich | Cat# G9023-102 |
| PFA 16% | Electron Microscopy Sciences | Cat# 15710-S |
| SDS | Sigma Aldrich | Cat# L3771-500G |
| Skin porcine gelatine | Sigma Aldrich | Cat# 48722-500G-F |
| Mowiol Mounting Medium | Homemade | - |
| Triton 10x | Sigma Aldrich | Cat# T9284-100ML |
| GentleMACS 25 C Tubes | Miltenyi Biotec | Cat# 130-093-237 |
| DNase I, RNase-free (1 U/μL) | Thermo Fisher Scientific | Cat# EN0521 |
| **Microscopes** |  |  |
| Clinostar 2 bioreactor | CelVivo |  |
| Echo Revolve2-K2-1861 microscope | Bico Groups |  |
| Leica SP8 confocal microscope | Leica Microsystems |  |
| **Software/ Python script** |  |  |
| Biorender | https://app.biorender.com/ |  |

## ACKNOWLEDGMENTS

CB was supported by Etat de Vaud, Swiss National Science Foundation (Grant Nos. 310030-182651 and 310030_215706), National Centre of Competence in Research Synapsy (Grant No. 51NF40-158776) (Switzerland), Novartis Foundation for Medical-Biological Research (Switzerland), Rub E. Funds (University of Lausanne, Switzerland), ERA-NET NEURON Joint Transnational Research Projects on Sensory Disorders (Grant No. 2020-088) (European Commision), Research Projects of Relevant National Interest (Grant No. 201789LFKB), Ministry of University and Research, Telethon Foundation (Grant No. GGP20137) (Italy), Opening the Future Program (KU Leuven), UCB Award, and Queen Elizabeth Foundation (Belgium). GA was supported by CelVivo Aps. We thank Prof. Peng Jin (Emory University) for the hiPSC lines. We acknowledge the Lausanne Genomic Technologies Facility (GTF) for single cell RNA sequencing and the Cellular Imaging Facility (CIF) for support.

## AUTHOR CONTRIBUTIONS

GA and CB designed the study and wrote the manuscript with inputs from all the authors. GA with help from MN, BV, CR performed all the experiments. AR designed the python-based script for morphological analysis and CR analysed the data. GA performed scRNAseq analyses. All co-authors provided feedback on the manuscript.

## SUPPLEMENTARY MATERIAL

**Dataset EV1.** Morphological features across conditions and timepoints.

**Dataset EV2.** Comparison of the morphological parameters’ variability.

**Dataset EV3.** MAD for each morphological parameter, comparing inter-batch variability between PA and RC conditions at day 6.

**Dataset EV4.** MAD for each morphological parameter, comparing inter-batch variability between OS and RC conditions at day 90.

**Movie EV1**. Particle trajectories and surface velocity magnitude.

**Movie EV2.** Organoids cultured in the RC apparatus.

## EXPANDED VIEW FIGURE LEGENDS

**Fig EV1.**
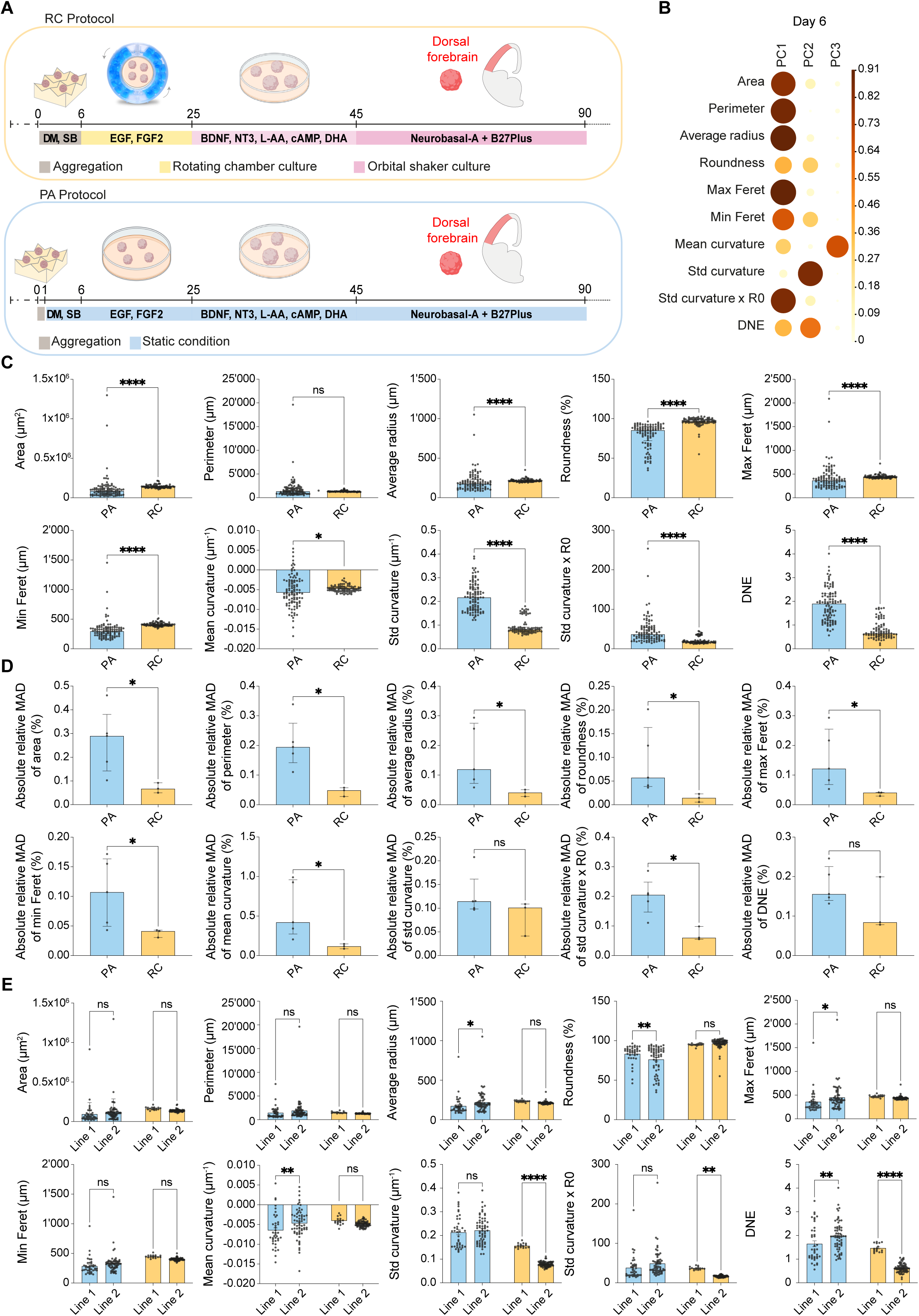
Prolonged aggregation time enhances EBs morphological homogeneity. **(A)** Schematic representation of the RC and PA protocols, highlighting key differences in culture conditions. RC organoids undergo a six-day aggregation phase followed by maintenance in a rotating chamber system, while PA organoids are aggregated for one day and cultured in static conditions. **(B)** Correlation plot showing the contribution of ten morphological parameters to the first three dimensions of the principal component analysis (PCA) at day 6. The importance of each parameter is represented as cosine² values across the principal components (related to Fig 2.E). The size and colour intensity of the circles are proportional to the parameters’ contribution to the principal components. **(C)** Bar plots showing quantitative comparisons of ten morphological parameters between PA (blue) (n = 102, five batches) and RC (yellow) (n = 88, three batches) EBs at day 6. Each dot represents an individual EB. The height of the columns represents the median and the error bars represent the interquartile range (i.e. the range between the 25^th^ percentile and the 75^th^ percentile). Statistically significant differences between conditions are indicated. Mann-Whitney test, ns = not significant, * p < 0.05, **** p < 0.0001. **(D)** Bar plots representing the absolute relative median absolute deviation (MAD) for each morphological parameter, comparing inter-batch variability between PA (blue) (n = five batches) and RC (yellow) (n = three batches) conditions at day 6. The height of the columns represents the median and each dot represents one batch, with error bars indicating interquartile ranges. Significant differences are marked. Mann-Whitney test, ns = not significant, * p < 0.05. **(E)** Bar plots depicting ten morphological parameters of EBs at day 6, analyzed across two independent iPSC lines (Line 1 and Line 2) cultured under PA (blue) (Line 1:n = 39, Line 2:n = 63) and RC (yellow) (Line 1:n = 16, Line 2:n = 72) protocols. Each dot represents an individual EB. The height of the bars represents the mean, and error bars indicate the standard error of the mean (SEM). Statistical significance was assessed using a two-way ANOVA with Šidák’s multiple comparisons test, with p-values adjusted for multiple comparisons. ns = not significant, * p < 0.05, ** p < 0.01, **** p < 0.0001.

**Fig EV2.**
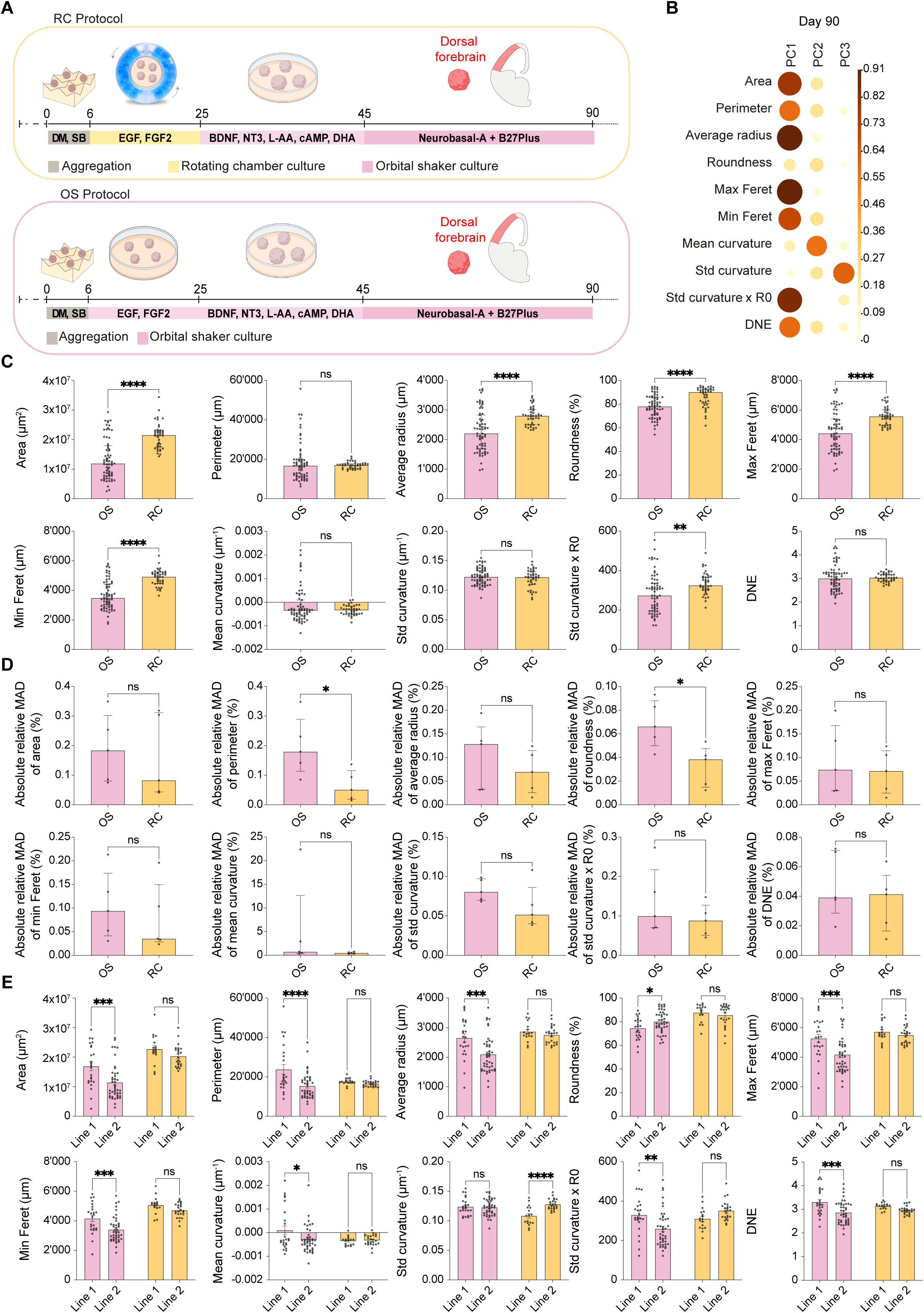
RC apparatus improves morphological homogeneity across batches and cell lines. **(A)** Schematic representation of the RC and OS protocols, highlighting key differences in culture conditions. From day 6 to 25, RC organoids are cultured in a rotating chamber, while OS organoids are maintained in a 10 cm plate on an orbital shaker **(B)** Correlation plot showing the contribution of ten morphological parameters to the first three dimensions of the principal component analysis (PCA) at day 90. The importance of each parameter is represented as cosine² values across the principal components (related to Fig 2.G). The size and color intensity of the circles are proportional to the parameters’ contribution to the principal components. **(C)** Bar plots quantifying ten morphological parameters in OS (pink, n = 67, five batches) and RC (yellow, n = 41, five batches) organoids at day 90. Each dot represents an individual organoid. The height of the bars represents the median, and the error bars correspond to the interquartile range (i.e., the range between the 25^th^ and 75^th^ percentiles). Statistically significant differences between conditions are indicated. Mann-Whitney test, ns = not significant, ** p < 0.01, **** p < 0.0001. **(D)** Bar plots showing the absolute relative median absolute deviation (MAD) for each morphological parameter, comparing inter-batch variability between OS (pink, n = 5 batches) and RC (yellow, n = 5 batches) conditions at day 90. The height of the columns represents the median and each dot represents one batch, and error bars indicate interquartile ranges. Significant differences are marked. Mann-Whitney test, ns = not significant, * p < 0.05. **(E)** Bar plots depicting ten morphological parameters of organoids at day 90, analyzed across two independent iPSC lines (Line 1 and Line 2) cultured under OS (pink) (Line 1:n = 24, Line 2:n = 43) and RC (yellow) (Line 1:n = 17, Line 2:n = 24) protocols. Each dot represents an individual organoid. The height of the bars represents the mean, and error bars indicate the standard error of the mean (SEM). Statistical significance was assessed using a two-way ANOVA with Šidák’s multiple comparisons test, with p-values adjusted for multiple comparisons. ns = not significant, * p < 0.05, ** p < 0.01, *** p < 0.001, **** p < 0.0001.

**Fig EV3.**
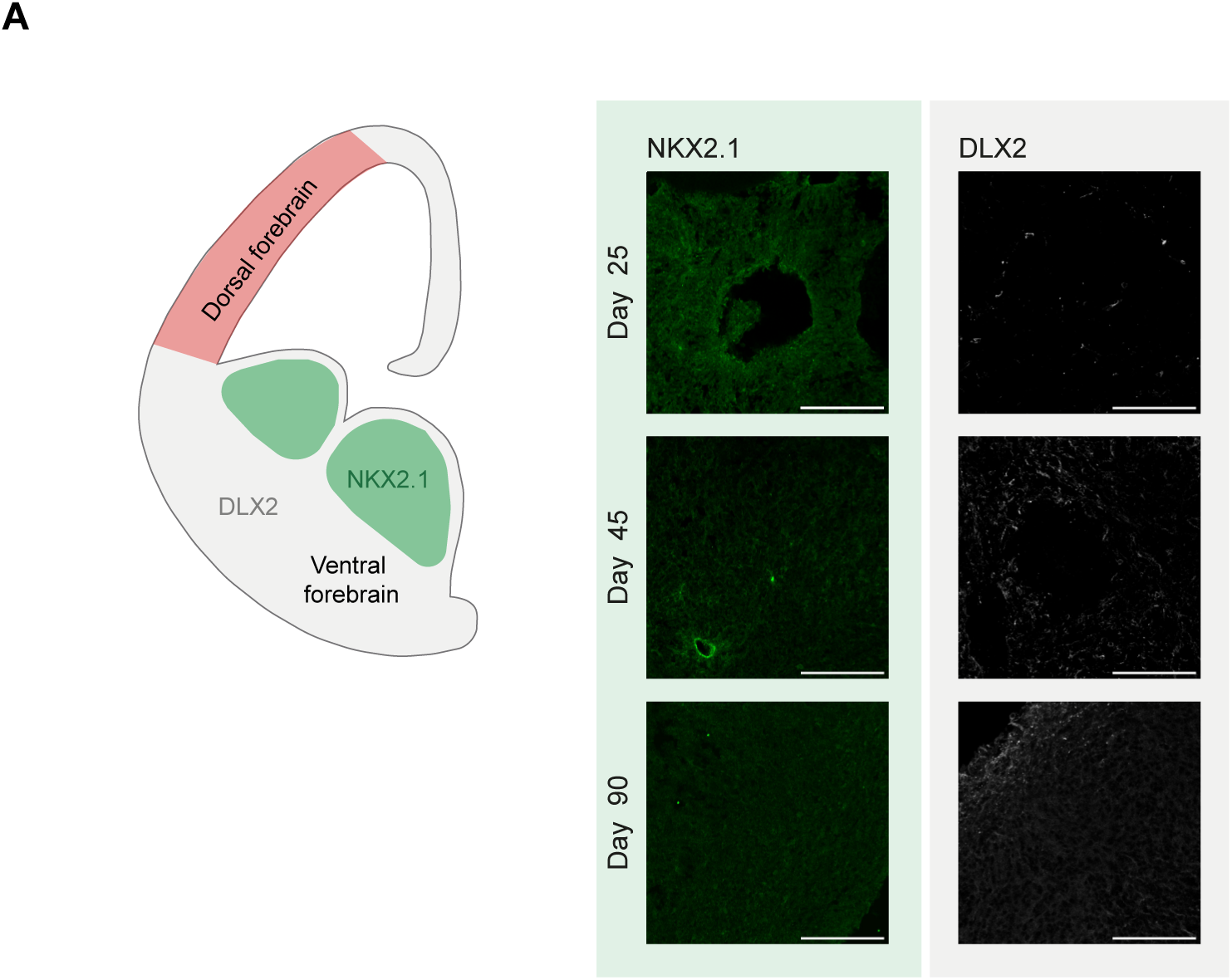
RC organoids do not show expression of ventral forebrain markers. **(A)** Schematic representation of the dorsal and ventral forebrain regions, highlighting the expression of ventral forebrain markers NKX2.1 and DLX2. **(B)** Immunofluorescence staining for NKX2.1 (green) and DLX2 (white) in RC-harvested organoids at days 25, 45, and 90. Scale bars: 100 μm.

**Fig EV4.**
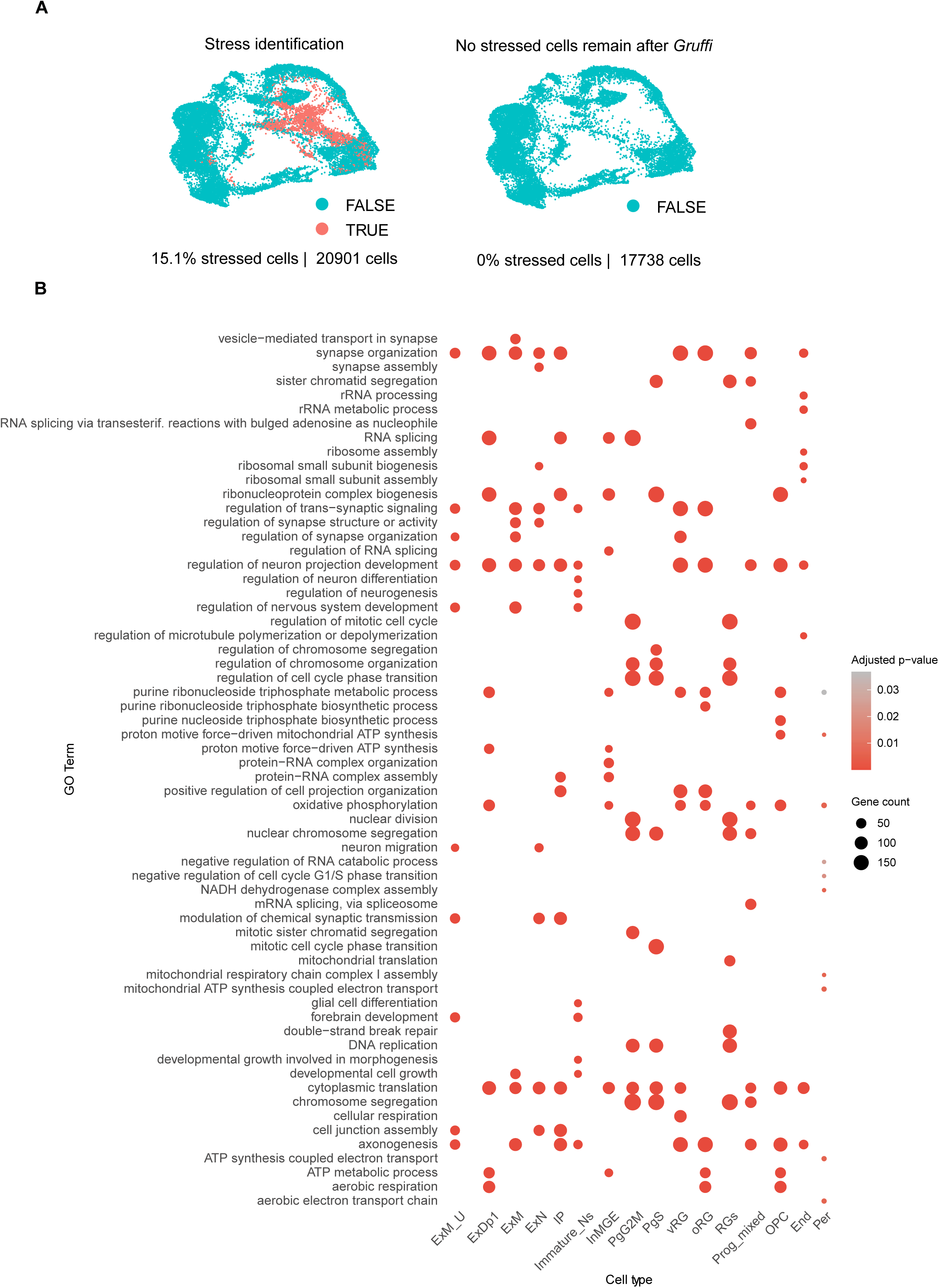
Quality control and functional annotation. **(A)** Identification and removal of stressed cells using the GRUFFI algorithm (Vertesy *et al*., 2022). The left UMAP plot displays the proportion of stressed cells (red) in the dataset before filtering, with 15.1% of cells (20,901 total) identified as stressed. The right UMAP plot shows the dataset after GRUFFI filtering, where all stressed cells have been removed, retaining 17,738 high-quality cells for downstream analysis. **(B)** Gene Ontology (GO) enrichment analysis of biological processes across different cell types in the dataset. The dot size corresponds to the number of genes enriched within each GO term, while the color intensity reflects the adjusted p-value, indicating the significance of enrichment. Cell types include: ExM_U (maturing excitatory upper enriched), ExDp1 (excitatory deep layer 1), ExM (maturing excitatory), ExN (migrating excitatory), IP (intermediate progenitors), Immature_Ns (immature neurons), InMGE (interneuron medial ganglionic eminence), InCGE (interneuron caudal ganglionic eminence), PgG2M (cycling progenitors in G2/M phase), PgS (cycling progenitors in S phase), vRG (ventricular radial glia), oRG (outer radial glia), RGs (radial glial populations), Prog_mixed (mixed progenitors), OPC (oligodendrocyte progenitor cells), End (Endothelial cells) and Per (pericytes).

